# High precision CA-ID-TIMS U-Pb zircon age for the Dueling Dinosaur locality, with implications for regional correlation, basal age and duration of the Hell Creek Formation, Montana

**DOI:** 10.1101/2025.07.10.664044

**Authors:** Eric M. Roberts, Marc S. Hendrix, Jahandar Ramezani, William C. Clyde, Pierre Zippi, Stuart Hodgson, Valerie Yuleridge, Lindsay E. Zanno

## Abstract

Discovery of the spectacular ‘Dueling Dinosaurs’ and other significant dinosaur localities from remote and isolated exposures of the Hell Creek Formation in central Montana highlight the complexity of establishing stratigraphic context and correlating Hell Creek Formation fossil localities located outside of the type area. This is particularly problematic for the lower two-thirds of the formation, which generally lacks reliable biostratigraphic or magnetostratigraphic zonation and has no dated ash beds. To address these enduring issues for one of the most significant Upper Cretaceous terrestrial fossil-bearing units in North America, detailed stratigraphic sections were established on the Murray Ranch and on McGinnis Butte in central Montana and correlated with other published Hell Creek Formation localities via magnetostratigraphy, biostratigraphy, and radioisotopic dating of ash beds. Results indicate that the K-Pg boundary is not exposed in the study area, however high-precision U-Pb CA-TIMS zircon ages for two newly discovered ash beds (66.929 ± 0.020 Ma and 66.850 ± 0.026 Ma, 2σ internal uncertainties) bracketing the ‘Dueling Dinosaurs’ locality provide the first absolute ages for the lower portion of the Hell Creek Formation, anywhere. Bayesian age-stratigraphic modelling places the ‘Dueling Dinosaurs’ locality at 66.897 +0.023/-0.028 Ma and suggest that the age of the base of the formation is ∼67.102 +0.710/-0.173 Ma (or older) in the study area. Comparison of stratigraphic architecture within the study area with published sections in the type area suggests that named sandstone marker horizons used for lithostratigraphic and sequence stratigraphic correlation in the type area have limited utility for regional correlation and need to be used with caution.

## Introduction

The Hell Creek Formation is one of Earth’s most iconic terrestrial rock records because it contains a diverse, abundant, and well-preserved vertebrate fauna (especially dinosaurs) and flora that existed just prior to the Cretaceous/Paleogene (K-Pg) boundary. Studied for over a century [1], the richly fossiliferous strata of the Hell Creek Formation remains a focus of investigation by researchers attempting to understand the tempo and mode of evolution and the causes, timing and nature of extinction at the end of the Cretaceous [2–18]. Recent paleontological and geological investigations have focused on the stratigraphy of the uppermost Hell Creek Formation and the K-Pg boundary [19–21]. Sprain et al. [20–21] reported over 60 distinct tephra deposits over a 70 m interval spanning the uppermost Hell Creek Formation to the Tullock-Lebo Member contact in the Fort Union Formation south of Fort Peck Reservoir close to the type area. This has resulted in a high level of temporal and stratigraphic resolution for this interval based on biostratigraphy, magnetostratigraphy, chemostratigraphy and a series of high precision radioisotopic age dates on ash beds. Although the uppermost Hell Creek Formation is stratigraphically and temporally well-resolved, the lower two thirds of the formation is not and no radioisotopic ages have been reported.

This is surprising given these intervals are equally fossiliferous and represent one of the premier sources of Upper Cretaceous terrestrial vertebrates and plant macrofossils in North America [6,10]. Moreover, age estimates for the base of the formation and total duration of the formation vary widely, and remain a point of confusion [19, 22–26].

As greater attention is being devoted to establishing a high-resolution geochronology throughout the Campanian-Maastrichtian of the Western Interior Basin [20–21, 27–35], the middle to lower portions of the Hell Creek Formation represents a critical stratigraphic interval in need of better documentation. This is particularly the case across much of the Hell Creek Formation in central and southern Montana, Wyoming and South Dakota. Most stratigraphic studies of the formation have been focused in the Fort Peck Reservoir area north of Jordan, MT (lectostratotype area) and in portions of western-central North Dakota [6, 21, 36–37], with very few studies outside of these two areas.

In recent years, exposures of the Hell Creek Formation southwest of Jordan, Montana have drawn attention for the preservation and excavation of spectacular dinosaur discoveries made on private land. However, many dinosaur specimens recovered from private lands often lack the same level of stratigraphic and taphonomic context as those excavated in the more well studied exposures along Fort Peck Reservoir and in North Dakota, limiting their scientific value. Perhaps the most publicized fossil discovery of this nature is the “Dueling Dinosaurs” [38], a unique fossil preserving nearly complete, entwined skeletons of a eutyrannosaurian (NCSM 40000) and *Triceratops* (NCSM 40001) hypothesized in public forums to have died locked in combat. NCSM 40000 and 40001 remained unavailable for scientific study until 2024, when they were officially accessioned at the North Carolina Museum of Natural Sciences (NCSM).

NCSM 40000 and 40001 were collected over two decades ago as part of commercial excavation. When gifted to the NCSM many details about the specimens’ stratigraphic and sedimentological context were limited. As a first step towards contextualizing this important locality and encouraging similar investigations of other sites with limited geological context, we returned to the original excavation site of the Dueling Dinosaurs (NCSM 40000 and 40001) and nearby McGinnis Butte with permission of the landowners and original collectors to establish a robust Hell Creek Formation reference section calibrated by high precision chronostratigraphy (Fig 1). Key goals of this study were to identify datable volcanic tephra beds or bentonites from the lower two thirds of the Hell Creek Formation and supporting magnetostratigraphic and palynologic data to better resolve the age of the fossils, to establish constraints on the total duration of the Hell Creek Formation in this area (and regionally), and to better document the stratigraphic relations between the Hell Creek Formation at this locality and correlative sections and units in the US and Canada.

**Fig 1.**
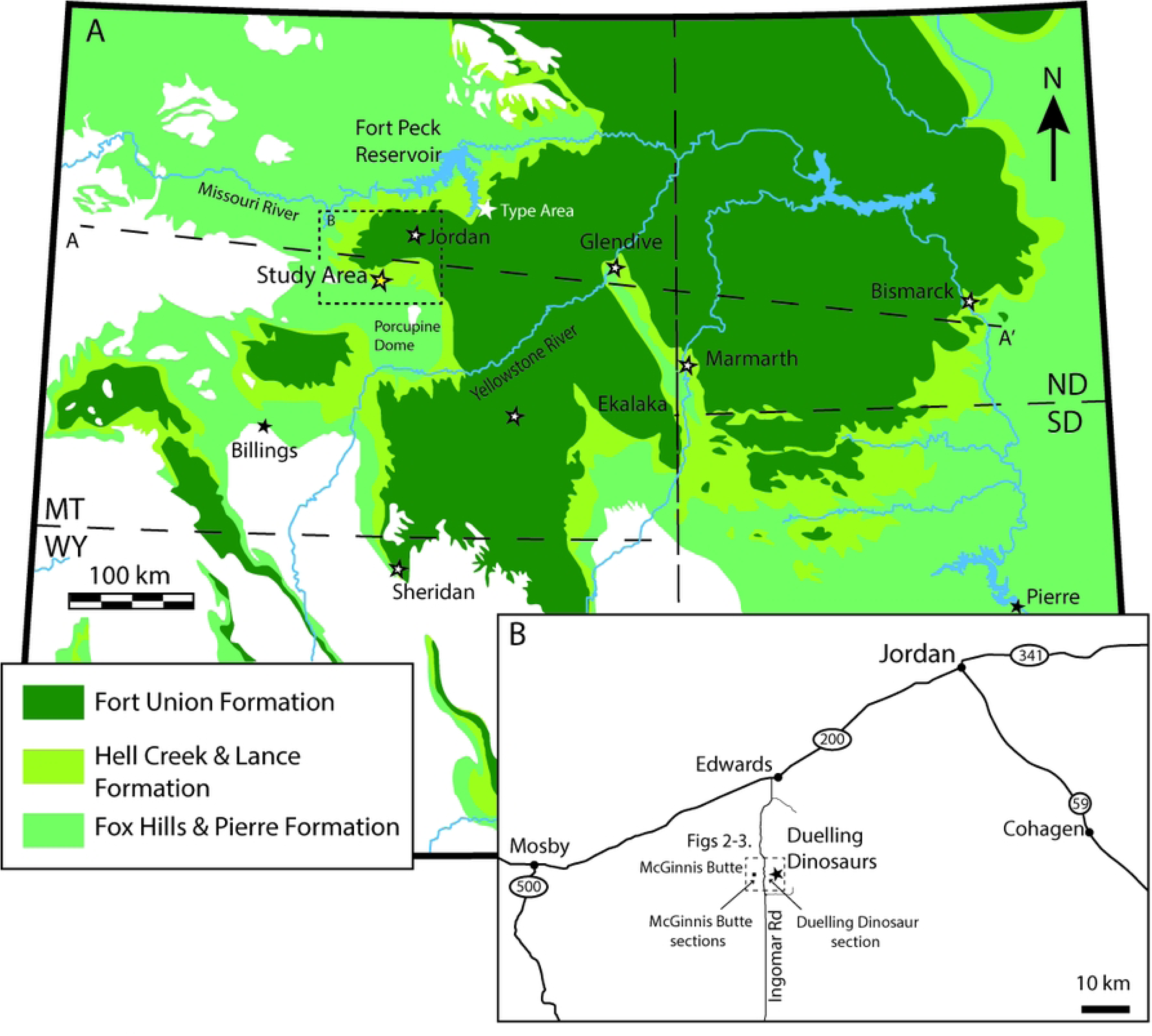
(A) Outcrop distribution of the Hell Creek Formation and bracketing formations across parts of Montana, North Dakota, South Dakota and Wyoming with location of the Dueling Dinosaurs locality and study area. (B) Detailed location map of the Dueling Dinosaur site and key measured sections within the study area.

## Geologic background

The Hell Creek Formation is an exceptionally well-exposed and continuous Upper Cretaceous (Maastrichtian) formation that crops out over parts of central to eastern Montana as well as western North and South Dakota (Fig 1). Initial study of the Hell Creek Formation was undertaken by paleontologist Barnum Brown between 1902 to 1909. Brown [1] identified a series of sandstone and mudstone units above the underlying Fox Hills Formation and below the overlying lignitic beds of the Fort Union Formation. Later work by Cobban and Reeside [39] elevated these beds to the formation level. More recent studies of the Hell Creek Formation have tended to divide the formation into the informal lower, middle, and upper Hell Creek designations [10–11, 26], although Frye [40] and others have suggested as many as seven members. Despite an apparent convergence towards a regional stratigraphic understanding of the Hell Creek Formation in more recent works, the formation is infamous for the complexity of its lateral facies relationships, with many stratigraphic units pinching out laterally and interfingering with facies that represent different sub-environments [7, 41]. Indeed, many workers have noted a paucity of true marker horizons in the formation and that their stratigraphic observations could not be correlated regionally except when either the upper or lower boundary of the formation could be observed [6, 11, 26]. Since Brown [1] did not formally define a stratotype section in his original description of the formation, the subsequent designation of a lectostratotype section at Flagg Butte (along a tributary of Hell Creek) by Hartman et al. [11] has alleviated some of these issues, particularly in and around the Fort Peck Lake region. However, it is unclear how comparable the Hell Creek Formation lectostratotype is outside of the Fort Peck region, and how well it applies to other fossiliferous sites that crop out tens or hundreds of kilometers away.

The Hell Creek Formation is also laterally equivalent (at least partially) to a number of other formations in the US and Canada, and the precise relationships remain untested in many cases, with limited information on the paleogeographic and paleoenvironmental variations between contemporary units. Key correlations include the Lance Formation in Wyoming and NE Colorado [6, 42], the Frenchmen and Scollard Formations in Saskatchewan and Alberta, Canada [25], the Denver Formation in eastern Colorado [43], the North Horn Formation in Utah [44], the Ojo Alamo and McRae formations in New Mexico [45–46] and the Javelina Formation in Texas [47].

The Hell Creek Formation is characterized by a strongly heterogenous series of interbedded sandstone and mudstone/siltstone units that are notably fossiliferous [1,11]. The contact with the overlying Fort Union Formation forms the Cretaceous/Paleogene (K-Pg) Boundary, an event associated with the extinction of all non-avian dinosaurs, as well as many other species of fauna and flora [2, 13, 42, 48–53]. Locally within the Hell Creek Formation, several marine tongues have been identified, including the Breien Member and Cantapeta Tongue, in the southeastern portions of the outcrop areas in North Dakota [7]. Otherwise, the Hell Creek Formation depositional environment has been interpreted as an extensive floodplain with meandering river systems and ephemeral backwater deposits [54]. The sandstone units are usually interpreted as large, extensive river systems [55], or channel complexes [26].

A significant amount of geochronological work has been conducted on the Hell Creek Formation, mostly focusing on the Hell Creek – Fort Union formational contact, due to its association with the K-Pg Boundary [42, 56]. The placement of the upper contact with the end-Cretaceous impact event (at least on a regional scale) constrains the upper age of the formation to ∼66.052 ± 0.008/0.043 Ma [20–21, 51], based on radiometric ages of numerous tephra-containing lignite beds common above and below the contact [20–21]. The age of the upper Hell Creek is also relatively constrained, thanks to an ash bed from an isolated coal seam (the Null Coal), which was dated via ^40^Ar/^39^Ar geochronology by Sprain et al. [20], yielding an age of 66.289 ± 0.051 Ma. However, no radioisotopic age determination has been conducted on strata below this level. As a result, age estimates for the base of the Hell Creek Formation, primarily based on magnetostratigraphy, are inconsistent ranging from 68.369 to 68.196 Ma [57], 67.47 to 67.5 Ma [37], 66.87 to 66.71 Ma [22]. Uncertainty surrounding sediment accumulation rates and the total duration of the Hell Creek Formation has, in part been attributed to the high degree of erosional scouring noted at the base of many of the fluvial sandstone units in the formation and apparent lateral variability between locations [26]. Collectively, stratigraphic and geochronologic uncertainties have led to differences in the estimated duration of deposition for the Hell Creek Formation that vary by >200%, ranging from 1.86 Myr [9], 1.36 Myr [22], 1.16 Myr [25], to 0.9 Myr [26].

## Methods

The focus of this investigation was to establish reference stratigraphic sections both at the Dueling Dinosaurs locality and nearby locations to build a well-constrained composite section in the region. Fieldwork was conducted at the Dueling Dinosaur site, as well as various other locations on the Murray Ranch and at the nearby McGinnis Butte section between 2018 and 2024. Three stratigraphic sections were hand-trenched and measured with a Jacob Staff. A full section through the study area was measured in two parts and called the lower and upper McGinnis Butte Sections (MB), and ∼2 km to the east, a partial section was measured through the Dueling Dinosaur Quarry, termed the Dueling Dinosaur (DD) section. This work was coupled with careful sampling of palynological, paleomagnetic, and volcanic tuff/bentonite samples for dating purposes.

## Palynology

Seventeen samples were collected for palynological investigation from hand-trenched sections (∼20-45 cm deep) to avoid surficial contamination and to obtain relatively unweathered samples. Sampling was focused on the upper parts of both the MB and DD sections to help locate the contact with the Fort Union Formation and the K-Pg boundary. After lithological examination and description, the samples were processed for palynology preparation. The samples were washed to remove surficial contaminants. Carbonate minerals were dissolved using HCl and silicate minerals removed using HF. After a wash with hot HCl, heavy liquid separation was performed with ZnBr_2_. The organic residue was washed with cold Schultz’s solution followed by a wash with ammonium hydroxide. The residues were sieved through a 7-μm mesh screen to remove small particles that would be unidentifiable in transmitted light microscopy. Organic residues were mounted on a coverslip with polyvinyl alcohol and fixed to a microscope slide with polyester resin. Slides were examined with phase contrast and differential interference contrast illumination using oil immersion at a minimum of 500X with a research grade Zeiss Axio Imager microscope. When possible, palynomorph occurrence data was collected until the total count reached 100 specimens for relative abundance data, after which, the remaining area of the slide was scanned for rare taxa that may have stratigraphic significance. Rare taxa were added to the count data. Key specimens were photographed.

## Magnetostratigraphy

Oriented paleomagnetic hand samples were collected from indurated fine-grained units by removing cover to expose fresh, relatively unweathered surfaces. Due to the poorly cemented, friable nature of many of the samples, we were not always able to collect replicate samples, and in some cases returned to the site to collect additional samples. Sampling strategy for paleomagnetic sampling was conducted following the acquisition of palynological analysis and radioisotopic ages of the two bentonite horizons, with the specific goal of locating the C30N/C29R and C29R/C29N reversals to establish stratigraphic control for the top of section in this area and potentially identify the bounds of Chron C29R that contains the K-Pg boundary. Stratigraphic spacing between sample sites varied but was typically between <1 m to 5 m through the primary interval of interest due to variability in the availability of appropriate lithologies. Five stratigraphic levels were sampled through the upper DD section, ranging from the DD quarry to the top of the local section, and ten stratigraphic levels were sampled from through the MB Section.

Samples from each location were cut into 8 cm^3^ cubes, retaining the oriented surface as one cube face, and analyzed at the University of New Hampshire Paleomagnetism Laboratory. Samples were measured using a 2G Enterprises superconducting quantum interference device (SQUID) cryogenic magnetometer shielded from the background magnetic field and step-wise demagnetized using a combination of an ASC Scientific Model TD-48SC thermal demagnetizer and AF Molspin tumbling alternating field (AF) demagnetizer. A mixed thermal and AF demagnetization protocol is consistent with the many previous paleomagnetic studies of the Hell Creek Formation which suggest titanomagnetite as the dominant NRM carrier [3–4, 19, 22, 21, 37] with intermediate titanohematite and goethite as potentially important ancillary magnetic carrier phases [58].

All sample data were analyzed using the PuffinPlot paleomagnetic data program [59]. Samples with three or more sequential steps exhibiting linear or quasi-linear decay to the origin were characterized using principal component analysis (PCA; [60]). Only those samples with a maximum angular deviation of ≤ 22° were included. A Fisher mean [61] was calculated for samples displaying an initial decay followed by strong clustering of vector endpoints and no further decay. Some samples with overlapping unblocking spectra display a demagnetization path best characterized by a great circle. Some samples exhibited unstable demagnetization behaviors and were excluded from further analysis.

## Bentonite sampling and U-Pb geochronology

Identification, collection, and tracing of two distinct bentonite units (devitrified volcanic ash beds), referred to herein as the Dueling and Ingomar bentonite beds, was carefully conducted to identify locations where the beds, which pinch in and out, were each purest. The Dueling and Ingomar bentonite beds are ∼5 m apart stratigraphically and can both be traced through the outcrop area; however, the two beds were sampled in separate locations ∼2 km apart from each other (Fig 2). This was done by digging large pits and excavating down onto the bentonite surface to collect fresh, uncontaminated samples (4-5 kg of material) from several mm above the basal surface of the bentonite (to avoid contamination with the underlying unit). Samples were processed in the lab by soaking in water for 1-2 days, followed by liquefaction using a blender and gradual clay disaggregation via a sonic dismembrator [62]. After reducing the sample to sand and silt sized grains (mostly volcanic phenocrysts), paramagnetic minerals were removed using a Franz magnetic separator, followed by a high-density liquid separation using methylene iodide to produce the heavy mineral separate. Final zircon selection was carried out by hand picking under a binocular microscope in which preference was given to sharply faceted, acicular zircon that contained elongate glass (melt) inclusions parallel to their long axis. Ramezani et al. [31] demonstrated in other Cretaceous Western Interior bentonite samples that these grains tend to yield the youngest eruption phase ages.

**Fig 2.**
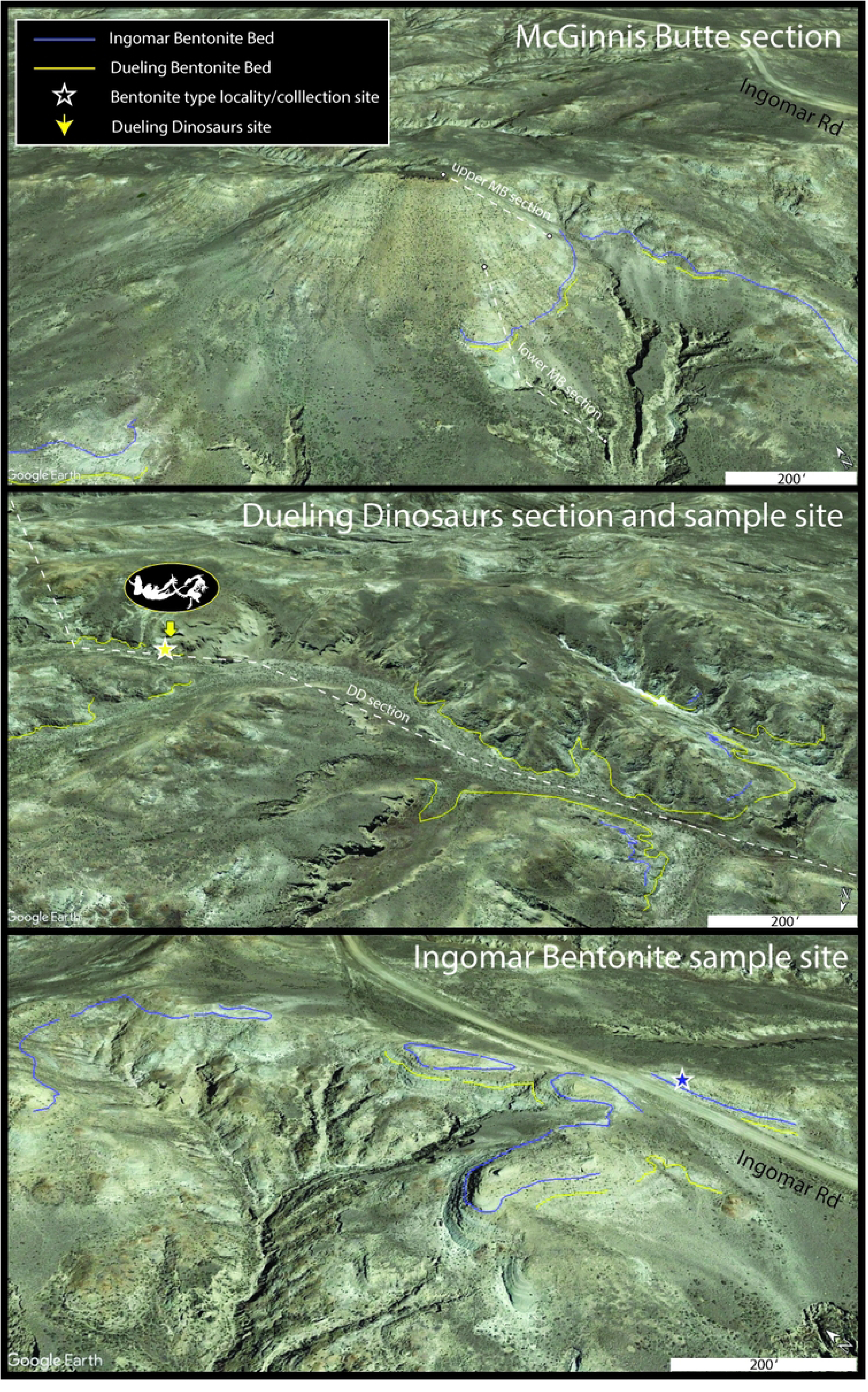
Locations of the Dueling and Ingomar Bentonites (imagery from GoogleEarth) (see Fig 1 for details). (A) McGinnis Butte with location of both lower and upper MB sections shown by dashed white lines and distribution of the Ingomar and Dueling bentonite beds shown by blue and yellow lines, respectively. (B) Dueling Dinosaur fossil (NCSM 40000 and NCSM 40001) location and sample location of the Dueling bentonite shown by yellow star. Dashed line shows trace of DD section (note the base and top of section not shown) and the distribution of the Dueling and Ingomar bentonite beds shown by yellow and blue lines, respectively. (C) Ingomar Bentonite sample location near Ingomar Road shown by purple star. Purple and yellow lines show the distribution of the two bentonites.

Zircons selected for U-Pb dating were pre-treated by a chemical abrasion technique modified after Mattinson [63], which involved thermal annealing in a 900°C furnace for 60 hours, followed by partial dissolution (leaching) in concentrated hydrofluoric acid (HF) to mitigate the effects of Pb-loss in zircon that result in anomalously young dates [31]. For leaching, annealed zircons were loaded with ∼75 µl of 29 M HF into 200 µl FEP Teflon® microcapsules, placed within a high-pressure Parr® vessel and left in a 210°C oven for 12-13 hours. This is considered a rather aggressive leach schedule but proven necessary as a remedy for persistent Pb loss in certain zircons (Widmann et al., 2019). The leached grains were transferred into 3 ml Savillex® FEP beakers and fluxed in successive steps in a dilute HNO_3_ solution and in 6N HCl over a hot plate (1 hour per step), with each step followed by agitation in an ultrasonic bath (1 hour) and rinsing with several milliliters of ultra-pure water to remove the leachates. Thoroughly rinsed zircon grains were loaded back into their microcapsules, spiked with the EARTHTIME ET2535 mixed ^202^Pb-^205^Pb-^233^U-^235^U isotopic tracer [64–65] and dissolved completely in 29M HF at 210°C for 48 hours.

Dissolved Pb and U were chemically separated using a miniaturized HCl-based ion-exchange chemical procedure modified after Krogh [66], using 50 µl columns of 1×8 anion-exchange resin. Purified Pb and U were loaded with a silica gel – H_3_PO_4_ emitter solution [67] onto single, degassed Re filaments and their isotopic ratios were measured on the Isotopx X62 multi-collector thermal ionization mass spectrometer equipped with a Daly photomultiplier ion counting system at MIT. Pb isotopic measurements were made on monoatomic Pb ions in a peak-hopping mode on the ion counter, whereas U isotopes were measured as UO ^+^ in a static mode on three Faraday detectors simultaneously.

Five zircons each were analyzed from the two bentonite beds in the Hell Creek Study Area. Data reduction, as well as calculation of U-Pb dates and propagation of uncertainties were accomplished using the Tripoli and ET_Redux applications [68–69]. Measured isotopic ratios were corrected for mass-dependent isotope fractionation in the mass spectrometer using the tracer ^202^Pb/^205^Pb and ^233^U/^235^U isotopic ratios, as well as for U and Pb contributions from the spike and laboratory blanks. Common Pb in the analyses averaged 0.37 pg, all of which was attributed to laboratory blank, and its isotopic composition was determined from long-term measurements of the total procedural Pb blank in the lab (see Table DR1 footnotes). The radiogenic ^206^Pb concentrations were also corrected for initial ^230^Th disequilibrium in magma using a magma initial Th/U model ratio of 2.8 ± 1.0 (2σ). This range of Th/U ratios encompasses all likely compositions of the magma source of an intermediate to felsic tuff [e.g. 70]. Pb isotopic ratios were corrected for isobaric interferences from Tl and BaPO_2_ on mass 205 by monitoring masses 203 and 201, respectively, and using natural isotopic abundances of ^138^Ba and ^205^Tl. Measured U isotopic ratios were also corrected for isobaric interference of ^233^U^18^O^16^O with ^235^U^16^O^16^O using an ^18^O/^16^O ratio of 0.00205 ± 0.00004 (2σ), which has been determined from long-term measurements of 272/270 mass ratio from large U loads. The present-day natural U isotopic composition of 137.818 ± 0.044 (2σ) was used in data reduction following [71].

In general, ^206^Pb/^238^U dates are considered the most precise and accurate in high-precision U-Pb geochronology as they are independent of suspected inaccuracy of the ^235^U decay constant [72–73], which potentially biases the ^207^Pb/^235^U or ^207^Pb/^206^Pb dates. Sample ages are derived from the weighted mean ^206^Pb/^238^U date of the five analyzed zircons in each sample, which form statistically coherent clusters. No analyses were rejected. Uncertainties in calculated ^206^Pb/^238^U dates are reported at 95% confidence level and in the ±*X*/*Y*/*Z* Ma format, where *X* is the internal (analytical) uncertainty in the absence of all external errors, *Y* incorporates *X* and the U-Pb tracer calibration errors and *Z* includes the latter as well as the decay constant errors of Jaffey et al. [74]. U-Pb bentonite ages presented here are discussed using the analytical uncertainty (X). When comparing these ages with data collected using different tracer solutions, analytical methods, or isotopic chronometers, consideration must be given to systematic contributions to uncertainty (i.e., Y and Z factors).

## Age-stratigraphic modelling

To construct a robust chronostratigraphic framework for the Hell Creek Formation at the Dueling Dinosaurs study area, we first established a composite stratigraphic section based on correlation of the DD and MB sections. To correlate the DD and MB sections, we walked out the Dueling and Ingomar bentonite beds between the sections, and we walked out the top contact of a prominent sandstone unit to correlate the Lower and Upper MB sections. We then employed a Bayesian age-stratigraphic model of the composite section generated using the Bchron software package [76–77]. The model utilizes the weighted mean dates of the two radiometrically-dated bentonites and a recalculated age for the C30n/C29r boundary based on high precision geochronology and magnetostratigraphy from the Denver Basin [27] modelled using the same Bayesian age-stratigraphic approach employed herein (66.48 +0.55/-0.17 Ma; larger of these asymmetric uncertainties was used in the DD-MB model) (see figure and data in Supplementary Materials). These input ages along with their relative stratigraphic positions were used to interpolate the age of Dueling Dinosaur site and the base of the formation. The underlying Markov chain Monte Carlo rejection algorithm of Bchron considers possible changes in the rock accumulation rate and results in more objective stratigraphic age uncertainties than the conventional linear extrapolation or spline-fit methods. The Bchron age model is shown with its median (solid) line and 95% confidence interval (shaded band). Code scripts, input data, and numerical model outputs are included in Supplementary Materials.

## Results

### Lithostratigraphy

Three stratigraphic sections were measured (lower and upper MB sections and DD Section; Figs 1 and 2) to establish a continuous composite stratigraphic framework through the study area. Sections were precisely correlated by tracing bentonite marker horizons between sections and walking out a continuous sandstone bench between the lower and upper MB sections (Figs 2 and 3). Although the underlying Fox Hills Formation was clearly identified in isolated locations throughout the study area, we are not entirely certain whether the proposed contact observed at the base of the Dueling Dinosaur Section and lower McGinnis Butte section represents the actual contact between Fox Hills and the Base of the Hell Creek Formation. The proposed top of the Fox Hills is marked by a major, multistory sandstone bench with persistent Fe-oxide staining and concretionary development, which was observed at the base of both sections in a poorly exposed sandstone unit at the base of dry stream beds in both sections. In nearby locations (within 0.5 km) where the Fox Hills Formation is better exposed, similar Fe-oxide staining and concretions are present, which lead us to this conclusion. However, the contact between the Fox Hills and Hell Creek formations has been described as highly variable from location to location [e.g., 26]. Complicating things further, we saw no evidence of the ‘toothpaste’ marker horizon in the study area or the Colgate Sandstone [79] in the area, which sometimes defines the Hell Creek-Fox Hills contact. Hence, we prefer to consider the Fe-Oxide-stained concretionary sandstone bench as the contact given that we see mostly covered interval below this level in the study area, with only isolated outcrops of Fox Hill Formation lower in section. However, we must consider the composite thickness of 109 m measured in the study area to be a minimum thickness for the Hell Creek Formation in the region (Fig 3).

**Fig 3.**
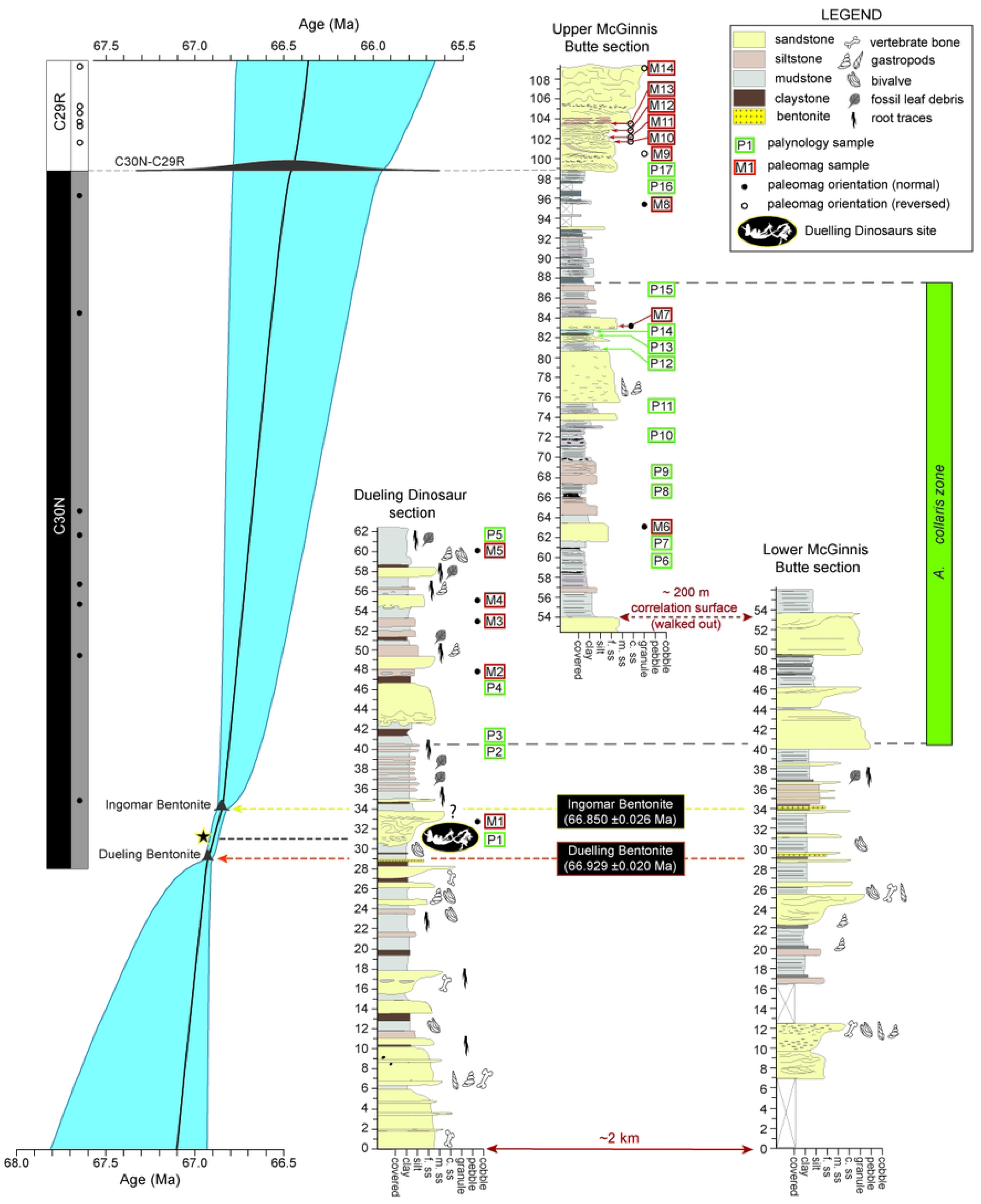
Correlated measured sections through the study area with accompanying chronostratigraphy and palynostratigraphy with Bayesian age-stratigraphic model on the left. Note that pollen and paleomag sample locations are shown by green and red boxes, respectively.

### Palynology

Palynomorph recovery and preservation ranged from marginal to excellent in the 13 samples analyzed, and a total of 107 individual species were identified with all species indicating a terrestrial or freshwater origin associated with fluvial, paludal, and lacustrine sub-environments (Table 1; Fig 4). Pollen diversity ranges from sample to sample. Some samples contain a low diversity palynoflora dominated by swamp fern spores and aquatic algae suggesting local source derivation and slow-moving or ponded water (e.g., samples P6, P7, and P8). Other samples contain a moderate or high diversity palynoflora that includes reworked mid-Cretaceous spores and dinocysts suggesting closer connection to a fluvial channel system (e.g., samples P9 and P13). A few samples were dominated by swamp fern spores suggesting floodplain deposition with reworked pollen indicating contributions from overbank flooding (e.g., sample P11). Except for some reworked middle Cretaceous marine dinoflagellates, no marine palynomorphs were encountered.

**Fig. 4.**
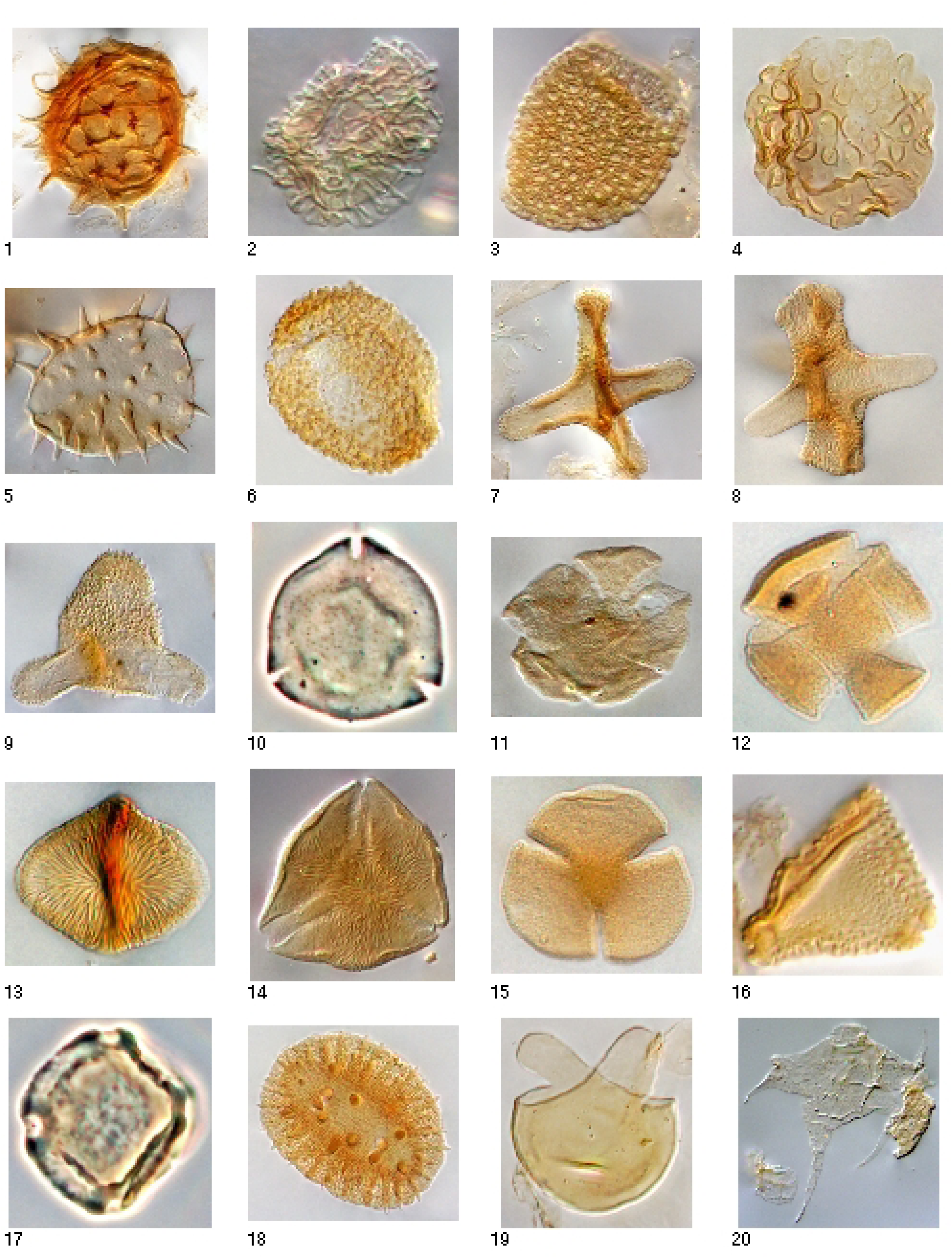
Photographs of diagnostic spores and palynomorphs documented in the Hell Creek Formation in the DD and MB sections. 1: *Ghoshispora bella*, 75 μm, P11 (9/94), L Maastrichtian-E Paleocene; 2: *Liliacidites altimurus* 26 μm, P3 (9.5/89), L Maastrichtian; 3: *Liliacidites complexus*, 40 μm, P5 (4/94), L Campanian-L Maastrichtian; 4: *Marsypiletes cretacea,* 44 μm, P1, Maastrichtian; 5: *Nuphar/Nypa/ Spinizonocolpites* sp., 45 μm, P3 (10/92.5), Latest Maastrichtian-Recent; 6: *Racemonocolpites formosus*, 52 μm, P9 (9/88), Latest Maastrichtian, except for one Latest Maastrichtian to Earliest Paleocene in Alberta; 7: *Aquilapollenites collaris*, 63μm, P11 (9/97), L Maastrichtian; 8: *Aquilapollenites conatus*, 54 μm, P3 (7/82.5), L Maastrichtian; 9: *Aquilapollenites delicatus*, 44 μm, 61822-4 10/98.5, L Maastrichtian; 10: *Kurtzipites circularis,* 23 μm, P9 (9/85), Maastrichtian-mid Paleocene; 11: *Leptopecopites pocockii*, 32 μm, P15 (12.5/76), Latest Maastrichtian; 12: *Scabrastephanocolpites lepidus*, p. 985, 25 μm, P9 (10/102), Campanian-Maastrichtian; 13: *Striatellipollis striatellus,* 28 μm, P8 (10/94), mid-L Maastrichtian; 14: *Styxpollenites calamitas*, 38 μm, P3 (9.5/89.5), Latest Maastrichtian; 15: *Tricolpites microreticulatus*, 24 μm, P7 (10/91), mid-L Maastrichtian; 16: *Tschudypollis retusus*, 23 μm, P11 (10/93.5), mid-Coniacian-L Maastrichtian; 17: *Ulmipollenites krempii*, 25 μm, P7 (10/92), L Maastrichtian-Recent; 18: *Wodehouseia spinata*, 50 μm, P1 (9/98.5), Latest Maastrichtian-Earliest Paleocene; 19: *Sigmopollis psilatus*, 29 μm, P11 (9/92), Maastrichtian-Recent; 20: *Nyktericysta davisii*, 95 μm, P2 (6.5/92), reworked from Middle-Late Albian marine strata.

**Table 1.** Palynological summary.

| Sample # | Strat. Section | Strat. Level (m) | Sample age | Sample age confidence | First Appearance (FA) species | FA species age | Last Appearance (LA) | LA species age |
| --- | --- | --- | --- | --- | --- | --- | --- | --- |
| P1 (21-DD-200) | DD | 30.6 | Latest Maastrichtian | high | *Ephedripites multipartitus* | Late Maastrichtian | *Aquilapollenites quadrilobus* | Latest Maastrichtian |
|  |  |  |  |  | *Marsipites cretacea* | Latest Maastrichtian | *Ephedripites multipartitus* | Late Maastrichtian |
|  |  |  |  |  | *Pandaniidites typicus* | Latest Maastrichtian | *Leptopocapites pocockii* | Late Maastrichtian |
|  |  |  |  |  | *Ulmipollenites krempi* | Late Maastrichtian | *Marsipites cretacea* | Latest Maastrichtian |
|  |  |  |  |  | *Ulmoidipites tricolatus* | Late Maastrichtian | *Tricolpites (Gunnera) microreticulatus* | Late Maastrichtian |
|  |  |  |  |  | *Wodehouseia spinata* | Latest Maastrichtian |  |  |
|  |  |  |  |  | *Ghoshispora bella* | Late Maastrichtian |  |  |
|  |  |  |  |  | *Triplanosporites sinuosus* | Late Maastrichtian |  |  |
| P2 (21-DD-203) | DD | 40.1 | Late Maastr-Paleocene | high | *Ulmipollenites krempi* | Late Maastrichtian | *Ulmipollenites krempi* | Paleocene |
|  |  |  |  |  | *Triplanosporites sinuosus* | Late Maastrichtian | *Cyathidites diaphana* | Paleocene |
|  |  |  |  |  |  |  | *Triplanosporites sinuosus* | Paleocene |
| P3 (61822-4) | DD | 42.1 | Latest Maastrichtian | high | *Ghoshispora bella* | Late Maastrichtian | *Osmundacidites?* | Late Maastrichtian |
|  |  |  |  |  | *Osmundacidites?* | Late Maastrichtian | *Aquilapollenites collaris* | Latest Maastrichtian |
|  |  |  |  |  | *Triplanosporites sinuosus* | Late Maastrichtian | *Aquilapollenites conatus* | Latest Maastrichtian |
|  |  |  |  |  | *Aquilapollenites collaris* | Latest Maastrichtian | *Aquilapollenites delicatus* | Late Maastrichtian |
|  |  |  |  |  | *Erdmanipollis cretaceus* | Late Maastrichtian | *Aquilapollenites quadrilobus* | Latest Maastrichtian |
|  |  |  |  |  | *Liliacidites altimurus* | Latest Maastrichtian | *Cranwellia rumseyensis* | Late Maastrichtian |
|  |  |  |  |  | *Styxpollenites calamitas* | Late Maastrichtian | *Liliacidites altimurus* | Latest Maastrichtian |
|  |  |  |  |  | *Ulmipollenites krempii* 4-pore | Late Maastrichtian | *Liliacidites complexus* | Latest Maastrichtian |
|  |  |  |  |  | *Ulmipollenites krempii* 3-pore | Late Maastrichtian | *Striatellipollis striatellus* | Late Maastrichtian |
|  |  |  |  |  | *Wodehouseia spinata* | Latest Maastrichtian | *Styxpollenites calamitas* | Latest Maastrichtian |
| P4 (61822-8) | DD | 46.8 | Latest Maastrichtian | high | *Osmundacidites?* | Late Maastrichtian | *Osmundacidites?* | Late Maastrichtian |
|  |  |  |  |  | *Triplanosporites sinuosus* | Late Maastrichtian | *Aquilapollenites collaris* | Latest Maastrichtian |
|  |  |  |  |  | *Aquilapollenites collaris* | Latest Maastrichtian |  |  |
|  |  |  |  |  | *Erdmanipollis cretaceus* | Late Maastrichtian |  |  |
|  |  |  |  |  | *Pandaniidites typicus* | Latest Maastrichtian |  |  |
|  |  |  |  |  | *Ulmipollenites krempii* 3-pore | Late Maastrichtian |  |  |
| P5 (61822-12) | DD | 61.8 | Latest Maastrichtian | high | *Liliacidites altimurus* | Latest Maastrichtian | *Liliacidites altimurus* | Latest Maastrichtian |
|  |  |  |  |  | *Pandaniidites typicus* | Latest Maastrichtian | *Liliacidites complexus* | Latest Maastrichtian |
|  |  |  |  |  | *Wodehouseia spinata* | Latest Maastrichtian | *Aquilapollenites quadrilobus* | Latest Maastrichtian |
| P6 (21-MB-35) | MB | 51.2 | Latest Maastrichtian | high | *Wodehouseia spinata* | Latest Maastrichtian | *Aquilapollenites delicatus* |  |
|  |  |  |  |  | *Ghoshispora bella* | Late Maastrichtian |  |  |
|  |  |  |  |  | *Triplanosporites sinuosus* | Late Maastrichtian |  |  |
| P7 (21-MB-03) | MB | 53.3 | Latest Maastrichtian | high | *Aquilapollenites collaris* | Latest Maastrichtian | *Aquilapollenites collaris* | Latest Maastrichtian |
|  |  |  |  |  | *Pandaniidites typicus* | Latest Maastrichtian | *Aquilapollenites quadrilobus* | Latest Maastrichtian |
|  |  |  |  |  | *Ulmipollenites krempi* | Late Maastrichtian | *Tricolpites (Gunnera) microreticulatus* |  |
|  |  |  |  |  | *Triplanosporites sinuosus* | Late Maastrichtian |  |  |
| P8 (21-MB-10) | MB | 58.7 | Late Maastrichtian | high | *Ephedripites multipartitus* | Late Maastrichtian | *Ephedripites multipartitus* | Late Maastrichtian |
|  |  |  |  |  | *Ulmipollenites krempi* | Late Maastrichtian | *Striatellipollis striatellus* | Late Maastrichtian |
|  |  |  |  |  | *Triplanosporites sinuosus* | Late Maastrichtian |  |  |
| P9 (21-MB-13) | MB | 60.85 | Latest Maastrichtian | high | *Cheno-Am* | Late Maastrichtian | *Aquilapollenites quadrilobus* | Latest Maastrichtian |
|  |  |  |  |  | *Ephedripites multipartitus* | Late Maastrichtian | *Ephedripites multipartitus* | Late Maastrichtian |
|  |  |  |  |  | *Erdmanipollis cretaceus* | Late Maastrichtian | *Tricolpites (Gunnera) microreticulatus* | Latest Maastrichtian |
|  |  |  |  |  | *Kurtzipites circularis/simplex* | Late Maastrichtian |  |  |
|  |  |  |  |  | *Pandaniidites typicus* | Latest Maastrichtian |  |  |
|  |  |  |  |  | *Racemonocolpites formosus* | Latest Maastrichtian |  |  |
|  |  |  |  |  | *Ulmipollenites krempi* | Latest Maastrichtian |  |  |
| P10 (21-MB-19) | MB | 64.8 | Latest Maastrichtian | high | *Aquilapollenites collaris* | Latest Maastrichtian | *Aquilapollenites collaris* | Latest Maastrichtian |
|  |  |  |  |  | *Erdmanipollis cretaceus* | Late Maastrichtian | *Aquilapollenites delicatus* | Late Maastrichtian |
|  |  |  |  |  | *Gaboniporis cristatus* | Late Maastrichtian | *Aquilapollenites quadrilobus* | Latest Maastrichtian |
|  |  |  |  |  | *Ulmipollenites krempi* | Late Maastrichtian | *Leptopocapites pocockii* | Late Maastrichtian |
|  |  |  |  |  | *Ghoshispora bella* | Late Maastrichtian |  |  |
|  |  |  |  |  | *Triplanosporites sinuosus* | Late Maastrichtian |  |  |
| P11 (21-MB-26) | MB | 67.7 | Latest Maastrichtian | high | *Alnipollenites trina* | Late Maastrichtian | *Aquilapollenites collaris* | Latest Maastrichtian |
|  |  |  |  |  | *Aquilapollenites collaris* | Latest Maastrichtian | *Aquilapollenites delicatus* | Late Maastrichtian |
|  |  |  |  |  | *Liliacidites altimurus* | Latest Maastrichtian | *Liliacidites altimurus* | Latest Maastrichtian |
|  |  |  |  |  | *Pseudoaquilapollenites conatus* | Latest Maastrichtian | *Pseudoaquilapollenites conatus* | Latest Maastrichtian |
|  |  |  |  |  | *Ulmipollenites krempi* | Late Maastrichtian | *Tricolpites (Gunnera) microreticulatus* | Late Maastrichtian |
|  |  |  |  |  | *Ghoshispora bella* | Late Maastrichtian | *Tschudyipollis (Proteacidites) retusus* | Late Maastrichtian |
|  |  |  |  |  | *Toroisporis major* | Late Maastrichtian | *Aquilapollenites quadrilobus* | Latest Maastrichtian |
|  |  |  |  |  | *Triplanosporites sinuosus* | Late Maastrichtian |  |  |
| P12 (21-MB-30) | MB | 73.7 | Late Maastrichtian | high | *Toroisporis major* | Late Maastrichtian | *Tricolpites (Gunnera) microreticulatus* | Late Maastrichtian |
|  |  |  |  |  | *Triplanosporites sinuosus* | Late Maastrichtian |  |  |
| P13 (21-MB-31) | MB | 74.2 | Latest Maastrichtian | high | *Alnipollenites trina* | Late Maastrichtian | *Aquilapollenites attenuatus* | Late Maastrichtian |
|  |  |  |  |  | *Aquilapollenites collaris* | Latest Maastrichtian | *Aquilapollenites collaris* | Latest Maastrichtian |
|  |  |  |  |  | *Ephedripites multipartitus* | Late Maastrichtian | *Ephedripites multipartitus* | Late Maastrichtian |
|  |  |  |  |  | *Erdmanipollis cretaceus* | Late Maastrichtian | *Liliacidites complexus* | Latest Maastrichtian |
|  |  |  |  |  | *Kurtzipites circularis/simplex* | Late Maastrichtian | *Pseudoaquilapollenites conatus* | Latest Maastrichtian |
|  |  |  |  |  | *Pandaniidites typicus* | Latest Maastrichtian | *Tricolpites (Gunnera) microreticulatus* | Late Maastrichtian |
|  |  |  |  |  | *Pseudoaquilapollenites conatus* | Latest Maastrichtian |  |  |
|  |  |  |  |  | *Ulmipollenites krempi* | Late Maastrichtian |  |  |
|  |  |  |  |  | *Ulmoidipites tricolatus* | Late Maastrichtian |  |  |
|  |  |  |  |  | *Triplanosporites sinuosus* | Late Maastrichtian |  |  |
| P14 (21-MB-33) | MB | 75.1 | Latest Maastrichtian | high | *Aquilapollenites collaris* | Latest Maastrichtian | *Aquilapollenites collaris* | Latest Maastrichtian |
|  |  |  |  |  | *Cheno-Am* | Late Maastrichtian | *Aquilapollenites quadrilobus* | Latest Maastrichtian |
|  |  |  |  |  | *Kurtzipites circularis/simplex* | Late Maastrichtian | *Tricolpites (Gunnera) microreticulatus* | Late Maastrichtian |
|  |  |  |  |  | *Ulmoidipites tricolatus* | Late Maastrichtian |  |  |
|  |  |  |  |  | *Ghoshispora bella* | Late Maastrichtian |  |  |
|  |  |  |  |  | *Triplanosporites sinuosus* | Late Maastrichtian |  |  |
| P15 (61822-1) | MB | 79 | Latest Maastrichtian | high | *Triplanosporites sinuosus* | Late Maastrichtian | *Aquilapollenites quadrilobus* | Latest Maastrichtian |
|  |  |  |  |  | *Pandaniidites typicus* | Latest Maastrichtian | *Aquilapollenites reticulatus* | Latest Maastrichtian |
|  |  |  |  |  | *Styxpollenites calamitas* | Late Maastrichtian | *Leptopocapites pocockii* | Late Maastrichtian |
|  |  |  |  |  | *Wodehouseia spinata* | Latest Maastrichtian | *Liliacidites complexus* | Latest Maastrichtian |
|  |  |  |  |  |  |  | *Styxpollenites calamitas* | Latest Maastrichtian |
|  |  |  |  |  |  |  | *Tricolpites (Gunnera) microreticulatus* | Late Maastrichtian |
| P16 (61722-4) | MB | 89 | Late Maastrichtian | high | *Sparganium* sp | Late Maastrichtian | *Aquilapollenites quadrilobus* | Latest Maastrichtian |
|  |  |  |  |  | *Ulmipollenites krempii* 4-pore | Late Maastrichtian | *Tricolpites (Gunnera) microreticulatus* | Late Maastrichtian |
|  |  |  |  |  | *Ulmipollenites krempii* 3-pore | Late Maastrichtian |  |  |
| P17 (61722-6) | MB | 90.3 | mid-late Maastrichtian | Low-moderate | *Tricolpites (Gunnera) microreticulatus* | Mid-Maastrichtian | *Tricolpites (Gunnera) microreticulatus* | Late Maastrichtian |

In spite of the various reworked material, all palynology samples from the Dueling Section and the Upper McGinnis Butte Section yielded late to latest Maastrichtian age estimates (Table 1), based on age ranges for palynomorphs reported from North America, with special emphasis placed on age ranges for palynomorphs reported from the Western Interior Seaway and the palynostratigraphy of Montana and surrounding areas more specifically [80–87]. No diagnostic Paleogene taxa were encountered in the highest samples collected ∼10 m below the top of the MB section. However, the distinctive biostratigraphic marker species *Aquilapollenites collaris* was observed to be common and first appears 40 m above the base of the DD/MB section or ∼9-10 m above the Dueling Dinosaur level. *A. collaris* is restricted to the middle and upper portions of the Hell Creek Formation and part of Subzone C of the *Wodehouseia spinata* Assemblage Zone [82]. The lack of this taxon in the lower 40 m of the section is consistent with this interval correlating to the lower portion of the Hell Creek Formation (Fig 3). Kaskes [37] in their stratigraphic investigation of the Naturalis *T. rex* locality, which is also located on the Murray Ranch ∼8 km to the east of the DD section, noted the first appearance of this taxon ∼10 m above the Naturalis *T. rex* locality. Interestingly, the first appearance of *A. collaris* in the DD/MB sections is also at ∼10 m above the Dueling Dinosaur fossil level, suggesting that the Dueling Dinosaurs and the Naturalis T. rex are likely both located at a similar stratigraphic level within the lower portion of the Hell Creek Formation [37]. Kaskes [3] placed the Naturalis site within the informal member nomenclature of Hartman et al. [2014], correlating the site with the top of the informal lower member. We avoid using this informal nomenclature as the sandstone beds used to define these boundaries cannot be confidently identified in the study area.

Along with palynology, kerogen analysis of the organic residue was also conducted on all samples. Charcoal is an abundant component of the kerogen in several samples, suggesting that forest fires were not uncommon in the area. Based on the thermal alteration index determined from kerogen, the Hell Creek Formation has undergone minor burial, with samples yielding estimated Ro% values between 0.25 and 0.34.

### Magnetostratigraphy

Most paleomagnetic samples exhibited stable demagnetization with relatively clear polarity determinations (Fig 5). All magnetostratigraphic samples collected from the DD section yielded normal polarity consistent with deposition during the C30n chron (Table 2). In contrast, a clear polarity reversal was documented between samples M8 and M9 within the MB section (Table 2; Fig 3). Approximately 5 meters of stratigraphy separates these two samples. Included in this interval is an erosional surface at the base of the ∼10 m thick sandstone that caps the section (Figs 2 and 3). We infer that the C30n/C29r reversal occurs at this erosional contact, which is ∼91.8 m level in the MB section (Fig 3).

**Fig 5.**
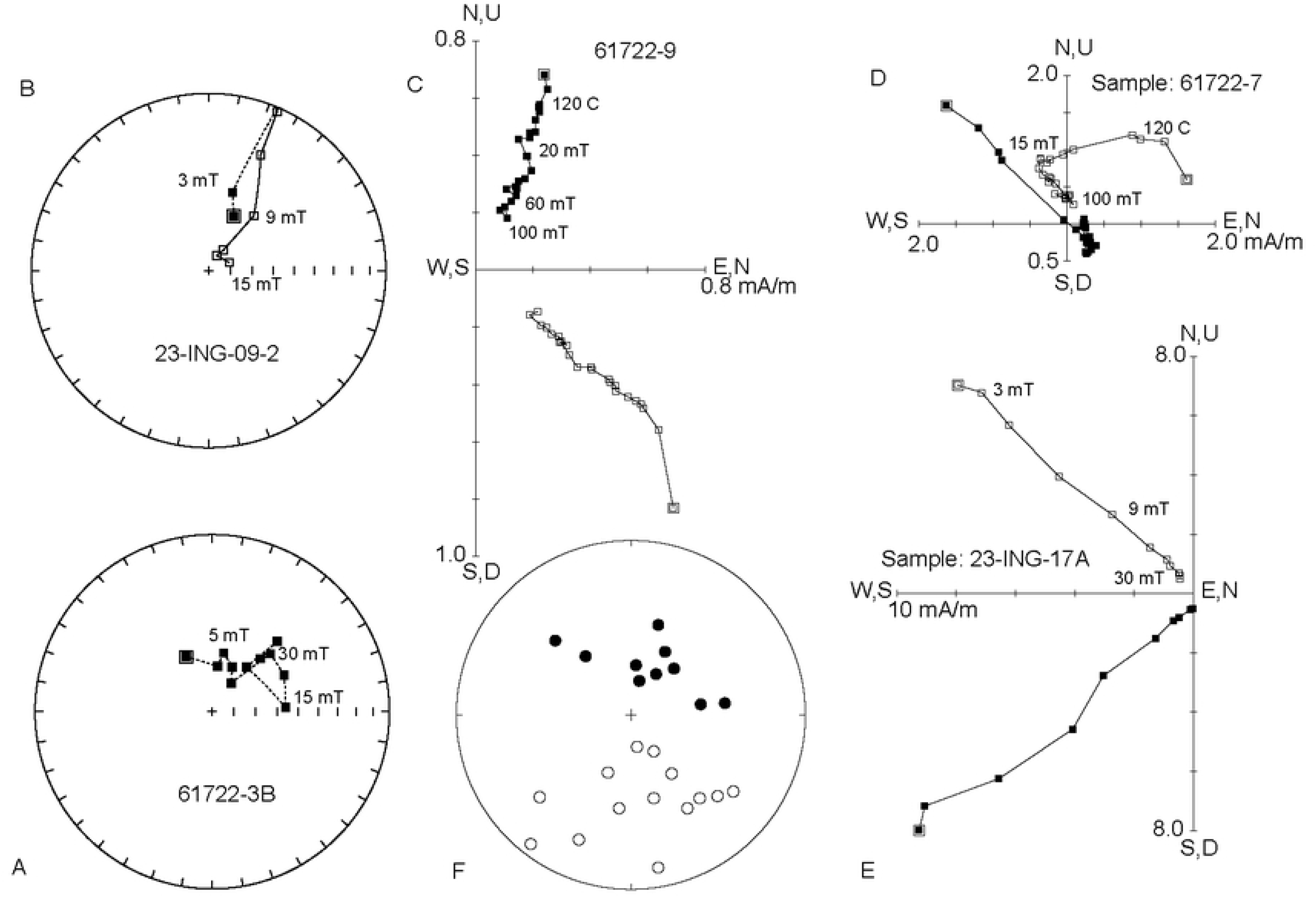
(A) Equal area projection of demagnetization data for a normal polarity sample illustrating clustering of remanence directions characterized by a Fisher mean [60]. (B) Equal area projection of demagnetization data for a reversed polarity sample where the remanence directions follow a great circle trajectory. (C-E) Vector endpoint diagrams for selected samples. Vector endpoints are labeled with alternating field intensity (mT) or temperature (°C). 61722-9 (normal polarity), 61722-7 (reversed polarity) and 23-ING-17A (reversed polarity) show linear decay toward the origin and are characterized by PCA. (F) Equal area projection of sample characteristic remanence directions for samples characterized by PCA or Fisher mean used for polarity interpretations.

**Table 2.** Summary of Paleomagnetic Data.

| Field Sample # | Sample # | Strat. Section | Strat. Level (m) | NRM intensity (A/m) | Total Steps | Dec. (deg) | Inc. (deg) | MAD/a95 | Steps in Calc | PCA/M/GC | Polarity |
| --- | --- | --- | --- | --- | --- | --- | --- | --- | --- | --- | --- |
| 61822-2 | M1 | DD | 33.2 | 2.07E-03 | 23 | 322.2 | 55 | 5.3 | 22 | PCA | N |
| 61822-7 | M2 | DD | 48 | 2.74E-04 | 17 | 13.5 | 73.8 | 18.5 | 11 | PCA | N |
| 23-DD-505B | M3 | DD | 53.7 | 1.90E-03 | 16 | 81.1 | 56.7 | 8.5 | 6 | M | N |
| 23-DD-505C-1 | M3 | DD | 53.7 | 3.43E-03 | 8 | 31.8 | 67.5 | 12.6 | 7 | PCA | N |
| 23-DD-505C-2 | M3 | DD | 53.7 | 2.48E-03 | 18 | 28.1 | 56 | 9.7 | 12 | PCA | N |
| 61822-9 | M4 | DD | 56.1 | 3.71E-04 | 23 | 5.9 | 66.8 | 10.3 | 22 | PCA | N |
| 61822-11 | M5 | DD | 60.8 | 4.04E-04 | 24 | 314.3 | 39 | 11.4 | 22 | PCA | N |
| 61722-9 | M6 | MB | 55.4 | 1.10E-03 | 23 | 16.7 | 45.2 | 10.4 | 23 | PCA | N |
| 61722-2 | M7 | MB | 75.8 | 6.79E-04 | 18 | 82.8 | 44.8 | 21.6 | 4 | M | N |
| 61722-3B | M8 | MB | 88 | 7.22E-05 | 12 | 43 | 60.1 | 8.4 | 11 | M | N |
| 23-ING-17A | M9 | MB | 93.4 | 1.39E-02 | 21 | 227.8 | -30.2 | 2.8 | 11 | PCA | R |
| 23-ING-17B | M9 | MB | 93.4 | 1.24E-02 | 27 | 170 | -12 | 1.8 | 11 | PCA | R |
| 23-ING-17C | M9 | MB | 93.4 | 2.45E-02 | 15 | 202.7 | -23.3 | 2.1 | 12 | PCA | R |
| 23-ING-16A | M10 | MB | 95 | 4.60E-03 | 13 | 148.1 | -69.8 | 9.7 | 9 | M | R |
| 23-ING-16B | M10 | MB | 95 | 3.54E-03 | 25 | 201.5 | -60.8 | 9.8 | 21 | M | R |
| 23-ING-301-B | M11 | MB | 95.4 | 9.64E-04 | 4 | 149 | -37.4 | 2.8 | 4 | M | R |
| 23-ING-301C | M11 | MB | 95.4 | 4.51E-02 | 18 | 133.2 | -32.2 | 16.2 | 4 | M | R |
| 23-ING-02 | M12 | MB | 96.4 | 1.90E-03 | 17 | 145.6 | -56.4 | 8 | 4 | M | R |
| 23-ING-09-1 | M12 | MB | 96.4 | 1.85E-03 | 9 | 187 | -45.2 | 14.6 | 3 | M | R |
| 23-ING-09-2 | M12 | MB | 96.4 | 1.39E-03 | 12 | 205.2 | 85.8 | 6.6 | 12 | GC | R |
| 23-ING-13 | M13 | MB | 96.7 | 3.96E-03 | 13 | 127 | -27.8 | 4.3 | 9 | PCA | R |
| 61722-7 | M13 | MB | 96.7 | 2.38E-03 | 25 | 165 | -48.9 | 14.2 | 14 | PCA | R |
| 1022-1A | M14 | MB | 101.7 | 2.61E-03 | 21 | 140.5 | -37.9 | 22 | 5 | PCA | R |
| 1022-1B | M14 | MB | 101.7 | 1.43E-03 | 13 | 221.3 | 82.8 | 14.2 | 10 | GC | R |
| 1022-2 | M14 | MB | 101.7 | 3.48E-02 | 15 | 169.1 | -74.8 | 4.4 | 9 | PCA | R |
| 1022-4 | M14 | MB | 101.7 | 5.57E-02 | 16 | 217.7 | -7.1 | 10.2 | 9 | PCA | R |
| Total Steps - total number of demagnetization steps |  |  |  |  |  |  |  |  |  |  |  |
| Dec/Inc for PCA line, Mean, or GC strike/dip of plane |  |  |  |  |  |  |  |  |  |  |  |
| MAD/a95 - MAD for PCA lines and GC, a95 for Means |  |  |  |  |  |  |  |  |  |  |  |
| Steps in Calc - Number of steps included in PCA, Mean, or GC calculation |  |  |  |  |  |  |  |  |  |  |  |
| PCA/M/GC - Sample characterized by Principle Component Analysis (PCA), Mean (M), or Great Circle (GC) |  |  |  |  |  |  |  |  |  |  |  |
| Polarity - Polarity determination based on PCA, Mean, GC direction |  |  |  |  |  |  |  |  |  |  |  |

### CA-ID-TIMS U-Pb geochronology and Bayesian model results

Two new U-Pb CA-ID-TIMS ages for the Hell Creek Formation from bentonite beds were successfully obtained from the study area. The Dueling Bentonite yielded an age of 66.929±0.020/0.021/0.076 Ma based on five concordant zircon analyses (Fig 6; Table 3). The stratigraphically higher Ingomar Bentonite (4.9 m above the Dueling Bentonite bed) yielded a slightly younger age of 66.850±0.027/0.026/0.076 Ma (Fig 6; Table 3). Both new ages, the recalculated age of the C30n/C29r boundary (66.48±0.55 Ma), and associated stratigraphic data were utilized to construct the Bayesian age-stratigraphic model shown in Fig 3. Unfortunately, the large uncertainty on the recalculated C30n/C29r chron boundary age and the unconstrained top of the section above this boundary (i.e., no K/Pg boundary present) means that the age model has limited utility for constraining the top of the section (see discussion). However, because the Dueling Dinosaur locality is bracketed between the two dated bentonites, a precise model age of 66.897+0.023/-0.028 Ma (asymmetric uncertainty) for the fossil bed was established (Fig. 3). Summary U-Pb analytical data is presented in Table 3 and in the Supplementary Material (S1 Table).

**Fig 6.**
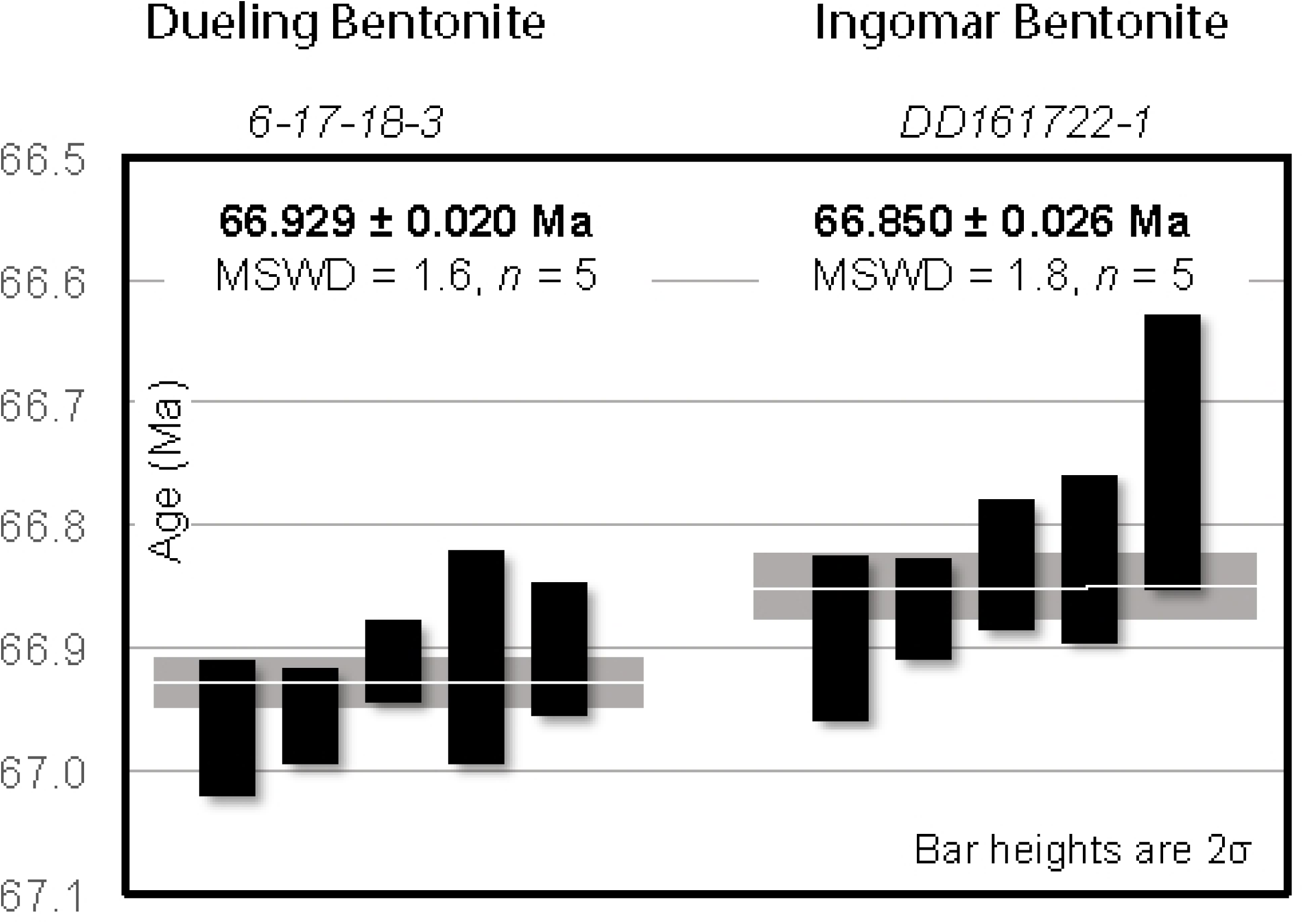
U-Pb Zircon concordia and weighted mean ages for the Dueling Bentonite and the Ingomar Bentonite.

**Table 3.** Summary of calculated U-Pb ages and their uncertainties.

|  |  |  |  |  | Ratios |  |  |  |  |  |  | Ages (Ma) |  |  |  |  |  |  |  |
| --- | --- | --- | --- | --- | --- | --- | --- | --- | --- | --- | --- | --- | --- | --- | --- | --- | --- | --- | --- |
| Sample | Pb(c) | Pb* | U | Th | 206 Pb | 208 Pb | 206 Pb |  | 207 Pb |  | 207 Pb |  | 206 Pb |  | 207 Pb |  | 207 Pb |  | corr. |
| Fractions | (pg) | Pb(c) | (pg) | U | 204 Pb | 206 Pb | 238 U | err | 235 U | err | 206 Pb | err | 238 U | err | 235 U | err | 206 Pb | err | coef. |
| (a) | (b) |  | (c) |  | (d) | (e) | (f) | (2σ%) | (f) | (2σ%) | (f) | (2σ%) |  | (2σ) |  | (2σ) |  | (2σ) |  |
| *Sample 6-17-18-3 (Dueling Bentonite)* |  |  |  |  |  |  |  |  |  |  |  |  |  |  |  |  |  |  |  |
| **z1** | 0.48 | 22.3 | 1008 | 0.42 | 1372.2 | 0.135 | 0.010442 | (.08) | 0.06861 | (.94) | 0.04768 | (.91) | **66.965** | **0.056** | 67.38 | 0.61 | 82 | 22 | 0.32 |
| **z4** | 0.33 | 36.2 | 1096 | 0.47 | 2192.1 | 0.149 | 0.010441 | (.06) | 0.06844 | (.60) | 0.04756 | (.58) | **66.956** | **0.039** | 67.22 | 0.39 | 77 | 14 | 0.34 |
| **z2** | 0.54 | 44.0 | 2218 | 0.49 | 2646.9 | 0.155 | 0.010434 | (.05) | 0.06823 | (.49) | 0.04745 | (.47) | **66.911** | **0.033** | 67.02 | 0.32 | 71 | 11 | 0.28 |
| **z5** | 0.55 | 12.5 | 649 | 0.44 | 777.0 | 0.140 | 0.010433 | (.13) | 0.06876 | (1.65) | 0.04782 | (1.61) | **66.908** | **0.087** | 67.5 | 1.1 | 89 | 38 | 0.32 |
| **z3** | 0.42 | 21.8 | 873 | 0.43 | 1343.8 | 0.136 | 0.010432 | (.08) | 0.06850 | (.95) | 0.04765 | (.93) | **66.901** | **0.053** | 67.28 | 0.62 | 81 | 22 | 0.32 |
| *Sample DD161722-1 (Ingomar Bentonite)* |  |  |  |  |  |  |  |  |  |  |  |  |  |  |  |  |  |  |  |
| **z3** | 0.29 | 20.3 | 541 | 0.50 | 1230.0 | 0.159 | 0.010431 | (.10) | 0.06855 | (1.17) | 0.04769 | (1.15) | **66.892** | **0.066** | 67.32 | 0.76 | 83 | 27 | 0.24 |
| **z2** | 0.26 | 55.0 | 1337 | 0.47 | 3312.9 | 0.152 | 0.010427 | (.06) | 0.06851 | (.60) | 0.04767 | (.58) | **66.868** | **0.041** | 67.28 | 0.39 | 82 | 14 | 0.42 |
| **z4** | 0.33 | 32.0 | 982 | 0.43 | 1959.2 | 0.139 | 0.010421 | (.08) | 0.06870 | (.70) | 0.04783 | (.69) | **66.832** | **0.053** | 67.47 | 0.46 | 90 | 16 | 0.24 |
| **z1** | 0.32 | 20.3 | 583 | 0.66 | 1175.3 | 0.210 | 0.010420 | (.10) | 0.06843 | (2.71) | 0.04765 | (2.69) | **66.827** | **0.068** | 67.2 | 1.8 | 81 | 64 | 0.17 |
| **z5** | 0.34 | 13.2 | 412 | 0.48 | 806.6 | 0.154 | 0.010407 | (.17) | 0.06896 | (2.53) | 0.04808 | (2.49) | **66.74** | **0.11** | 67.7 | 1.7 | 102 | 59 | 0.27 |
| **Notes:** |  |  |  |  |  |  |  |  |  |  |  |  |  |  |  |  |  |  |  |
| (a) Thermally annealed and pre-treated single zircon. Data used in age calculation are in bold. |  |  |  |  |  |  |  |  |  |  |  |  |  |  |  |  |  |  |  |
| (b) Total common-Pb in analyses. |  |  |  |  |  |  |  |  |  |  |  |  |  |  |  |  |  |  |  |
| (c) Total sample U content. |  |  |  |  |  |  |  |  |  |  |  |  |  |  |  |  |  |  |  |
| (d) Measured ratio corrected for spike and fractionation only. |  |  |  |  |  |  |  |  |  |  |  |  |  |  |  |  |  |  |  |
| (e) Radiogenic Pb ratio. |  |  |  |  |  |  |  |  |  |  |  |  |  |  |  |  |  |  |  |
| (f) Corrected for fractionation, spike and blank. Also corrected for initial Th/U disequilibrium using radiogenic <sup>208</sup>Pb and Th/U<sub>magma</sub> = 2.8. |  |  |  |  |  |  |  |  |  |  |  |  |  |  |  |  |  |  |  |
| All common Pb assumed to be laboratory blank. Total procedural blank less than 0.1 pg for U. |  |  |  |  |  |  |  |  |  |  |  |  |  |  |  |  |  |  |  |
| Blank isotopic composition: <sup>206</sup>Pb/<sup>204</sup>Pb = 18.15 ± 0.47, <sup>207</sup>Pb/<sup>204</sup>Pb = 15.30 ± 0.30, <sup>208</sup>Pb/<sup>204</sup>Pb = 37.11 ± 0.87. |  |  |  |  |  |  |  |  |  |  |  |  |  |  |  |  |  |  |  |
| Corr. coef. = correlation coefficient. |  |  |  |  |  |  |  |  |  |  |  |  |  |  |  |  |  |  |  |
| Ages calculated using the decay constants λ<sub>238</sub> = 1.55125E-10 y<sup>-1</sup> and λ<sub>235</sub> = 9.8485E-10 y<sup>-1</sup> (Jaffey et al. 1971). |  |  |  |  |  |  |  |  |  |  |  |  |  |  |  |  |  |  |  |

## Discussion

### Stratigraphy and age of the Dueling Dinosaur and McGinnis Butte sections

To date, the Dueling and Ingomar bentonites are the oldest dated tephra beds in the Hell Creek Formation. The Null Coal tephra (66.289 ± 0.051 Ma; [20]) is located several meters above the C30n/C29r chron boundary in the type area, whereas the Dueling and Ingomar bentonites occur ∼65-70 m below this chron boundary. In addition, these two bentonites occur 7 and 12 m below the first appearance of *A. collaris*, which is restricted to the informal middle member of the Hell Creek Formation elsewhere [82, 88]. Thus, the report herein of the Dueling and Ingomar bentonite ages and their stratigraphic position significantly expand the range of datable tephra deposits in the Hell Creek Formation while also demonstrating the potential for better age control throughout the formation.

Results of our Bayesian age-stratigraphic model indicate an age of 66.897+0.023/-0.028 Ma for the Dueling Dinosaur fossils and based on comparison of the pollen and stratigraphy of Kaskes [37] for the Naturalis *T. rex* site (∼8 km away), we conclude that both specimens are roughly the same age, located near the top of the informal lower member of the formation. Due to a lack of continuous exposure between the two sites, however, we cannot demonstrate whether both specimens occur within the same or different channelized sandstone complexes. Ongoing sedimentological investigation of channel sandstones in the study area suggests they have an anabranching morphology with multiple active channels at the same level (particularly in the lower half of the formation). For this reason and the likelihood that any aerially widespread channelized sandstone bodies are time-transgressive, we challenge the assumption that channelized sandstones provide a viable basis for lithostratigraphic or chronostratigraphic correlation in the Hell Creek Formation (e.g., JenRex Sandstone, Apex Sandstone, etc. [11, 26]. Our observations while walking out the Dueling bentonite in the study area reveal it to be highly discontinuous, primarily due to the anastomosed fluvial morphology in which multiple channels occur at the same stratigraphic level and separate broad packages of floodbasin mudrock (Fig 3; see also [89]).

### Age of base of Hell Creek Formation and its duration

The age-stratigraphic model produced in this study predicts an approximate age of 67.102 +0.710/-0.173 Ma for the base of the DD/MB composite section, which we propose as the base of the Hell Creek Formation in the study area. This model age has the advantage of being generated using the first dated ash beds from the lower portion of the formation, although we consider this a minimum age for the base of the Hell Creek Formation, regionally, since we cannot confidently confirm that the base of the section sits at the Fox Hills Formation contact in the study area. Furthermore, the precision of the basal model age is poor due to a sparsity of dated horizons close to the formation contact whereby variation in sediment accumulation rates below the Dueling bentonite could result in a younger or, especially, older true age for the contact. Comparison to other published estimates for the basal age of the Hell Creek Formation supports the age calculated here, which is essentially the same as the minimum age of 67.2 Ma suggested by Eberth and Kamo [25] based on their correlations with the Battle Formation in Alberta.

The C30n/C29r chron boundary age was crucial in generating the age-stratigraphic model herein, although determining the ideal age and uncertainty to use for this boundary was challenging. Sprain et al. [21] reported an age 66.311 ± 0.102 Ma (2σ analytical uncertainty) using linear extrapolation from ^40^Ar/^39^Ar geochronology of volcanic ash horizons close to the chron boundary in the Hell Creek Formation type area near Fort Peck Reservoir. Application of this age for our purposes would require the addition of systematic errors including decay constant uncertainty for both the boundary age and our dated bentonites, which would have a significant impact on the apparent precision of the model age for the Dueling fossil locality. Alternatively, Clyde et al. [27] calculated an age of 66.436 ±0.039 Ma (2σ analytical uncertainty) for the chron boundary as it occurs in the Denver Basin based on high-precision U-Pb zircon ages from volcanic ash horizons in the Kiowa core. This age is statistically indistinguishable from that of Sprain et al. [21] even before considering systematic error, although linear interpolation used by Clyde et al. [27] was applied across a much larger stratigraphic interval of ∼20 m with no additional age constraint below the chron boundary. As a result, the propagated uncertainty does not account for variability in sediment accumulation rates and is thus likely to significantly overestimate precision. To ensure the C30n/C29r chron boundary age was compatible with our DD/MB model, we recalculated the Clyde et al. [27] U-Pb zircon-derived age using a Bayesian age-stratigraphic model for the Kiowa core. Our recalculated chron boundary model age of 66.48+0.55/-0.17 Ma more appropriately represents stratigraphically propagated uncertainty but appears significantly less precise owing to a lack of dated horizons low in the Kiowa core. This issue is also present in the DD/MB model, raising concerns over the utility of both the recalculated chron age and the basal DD/MB model age. Future efforts to date volcanic ash horizons close to and below the C30n/C29r chron boundary would greatly improve temporal resolution of the Hell Creek Formation, as would a similar sampling approach above and below the basal contact of the formation.

Although the K-Pg boundary was not identified in our study area, we can estimate a total duration for the Hell Creek Formation in the study area to represent ∼1.08 Myr, based on our model age for the base of the DD/MB section (67.102 +0.710/-0.173) and the Clyde et al. [27] K-Pg boundary age of 66.021 ± 0.024 Ma (2σ analytical uncertainty), which is within error of the K-Pg age of Sprain et al. [21] and the GTS2020 [78]. The duration of deposition represented by the 109 m-thick DD/MB composite section (missing upper contact) is approximately 0.737 Myrs, equating to an average sediment accumulation rate of 25.8 cm/kyr across the 109-metre-thick section. Based on the time interval between top of the DD/MB section (model age of 66.365 +0.406/-0.713 Ma) and the Clyde et al. [27] K/Pg age of 66.02 Ma combined with the average sediment accumulation rate, the model predicts that ∼90 m of Hell Creek Formation strata may be missing above the MB section for a total formation thickness of ∼200 m in the study area at the time of deposition, not accounting for possible variation in sediment accumulation rates or unconformities. This would make the section considerably thicker than what has been reported in the type section and most other locations. Based on observations of Fort Union Formation cropping out several kms north of the study area, we suspect that 200 m is too thick of an estimate. However, the thickest sections of the Hell Creek/equivalent strata do reportedly thicken significantly to the south, even though minimal west-east thickening has been observed [90].

Using a previous K/Pg boundary age of 65.51 Ma, Hicks et al. [22] estimated the duration of the Hell Creek Formation at 1.36 Myr, however using the currently accepted boundary age of around 66.021 Ma [21], their estimate would be closer to a duration of between ∼700-850 kyr in western North Dakota. This is somewhat shorter in duration than what we report here, and may be explained by a possible eastward thinning, time-transgressive nature of the Hell Creek clastic wedge. It should be expected that the temporal duration of the Hell Creek Formation should increase to the west, and currently, the DD/MB Hell Creek Formation section is among the furthest west stratigraphic sections published for the formation, located south and slightly west of the type section at Flagg Butte near Reid Creek [11]. Although the stratigraphic relationships of the upper Hell Creek Formation are well resolved in the type area and well-studied sections in North Dakota, much work remains to understand the regional stratigraphic relationships of the Hell Creek Formation and correlative units, such as the Lance and St. Mary River formations, outside of these areas. Furthermore, until regional (i.e., basin wide) documentation of Hell Creek stratigraphic relationships are better resolved, fundamental uncertainties regarding the nature and timing of dinosaur (and other vertebrate) coevolution with their environment remain.

### Source of ash and correlation of bentonites within the Hell Creek Formation

Unlike most previous studies where tephras have primarily been identified within lignite seams in the Hell Creek and Fort Union Formation, we report two new thin, but pure bentonite beds from the lower portion of the Hell Creek Formation. These are the first two dated tephras documented from anywhere in the formation below the null coal ash (uppermost Hell Creek Formation). Their relative thinness and discontinuous but laterally reoccurring distribution, coupled with relatively fine phenocryst sizes (< 500 µm) suggests that the ashes are reasonably far travelled, or alternatively from less explosive, localized eruptions potentially emanating from nearby alkaline volcanic centers in the Central Montana Alkalic Belt. Campanian bentonites across the Western Interior Basin (WIB), from Alberta, Montana, Wyoming, Nebraska, and the Dakotas have been linked to the Elkhorn Mountain Volcanics [91–95]. However, the source of Maastrichtian ashes within the WIB remains far more speculative, with limited previous work.

As an example, the ages of the Dueling Bentonite and the Battle Bentonite in Alberta are within error of each other [25], leading us to speculate that both bentonites are part of the same ashfall event that may represent a regional marker horizon with a minimum area of 650 km^2^. Confirmation requires testing through geochemical fingerprinting, but it poses the question of just how extensive these ash beds are across western North America and whether they represent locally derived ashfall/ashflow deposits or large, regional tephra blankets from a single source. Given that so little is known about the nature and source of Maastrichtian (and Danian) bentonites in the Western Interior Basin, this presents a fertile area for future investigation. It is highly likely that with further exploration of the lower to middle portion of the Hell Creek Formation, additional ash beds will be discovered that will assist with this provenance interpretation, as well as lead to even better age control in the formation and establishment of regional marker horizons for correlation.

## Conclusions

Given the scientifically significant nature of the Dueling Dinosaur fossil locality, collected over a decade ago and recently accessioned into the North Carolina Museum of Natural Sciences collections, the primary aim of this study was to return to the Murray Ranch to establish detailed stratigraphic context for the “Dueling Dinosaurs” (NCSM 40000 and NCSM 40001). Due to the remote and poorly understood stratigraphy of the Hell Creek Formation in the extensive exposures in central Montana to the south and west of Jordan Montana, it was necessary to establish a composite section for the Murray Ranch and nearby McGinnis Butte by utilizing a variety of dating approaches to date and correlate the stratigraphy and significant vertebrate fossils, including NCSM 40000 and NCSM 40001. Lithostratigraphy, coupled with palynology and magnetostratigraphy, demonstrate that the composite section in the region is ∼109 m thick, beginning at or close to the Fox Hills Formation, and continuing to close to the top of the Hell Creek Formation, but ending just prior to the K-Pg boundary in lower C29R. Identification and high-precision CA-ID-TIMS U-Pb zircon dating of two bentonite beds bracketing the fossil locality are the stratigraphically lowest ever reported from the Hell Creek Formation. Based on these geochronologic results and Bayesian age-stratigraphic modelling, the Dueling Dinosaurs (NCSM 40000 and NCSM 4000) locality is assigned a precise model age of 66.895+0.023/-0.027 Ma, which roughly correlates with the top of the informal lower member. The two new U-Pb ages were also used to model a minimum age of 67.102 +0.710/-0.173 Ma for the base of the Hell Creek Formation the study area. This date is consistent with the age proposed for the base of the formation by Eberth and Kamo [25] and addresses the significant amount of uncertainty regarding the age of the base and total duration of the formation. Indeed, the CA-ID-TIMS U-Pb age of the Battle Bentonite (66.936 ±0.047 Ma) utilized by Eberth and Kamo [25] to make these estimates is within analytical uncertainty of the age of the Dueling Bentonite (66.929 ±0.020 Ma). It is thus likely that the Dueling/Battle bentonites represent the same ash bed forming a regional marker horizon with a minimum total areal extent of 650 km^2^. This presents a much-needed opportunity for correlation of the lower/mid Hell Creek Formation and equivalent strata, flora and fauna across the Western Interior Basin in the US and Canada. It is also important to point out that there are many such specimens collected by private collectors that have ended up in museums and that these specimens could also greatly benefit from follow-up geological investigations that have the potential to add important context to these specimens and can be achieved long after the original excavations took place.

## Acknowledgements

The authors wish to thank Mary Anne Murray for generous access to her property to study the geologic context of the site, and to Clayton Phipps for showing us the site and valuable discussions. We are also grateful to Eric Lund for field assistance. We thank Sarah Widlansky and Galen Walton for helping with processing and analyzing paleomagnetic samples. We also thank Russ Harms, Global Geolab Ltd for assisting with the pollen separations.

## SUPPLEMENTARY INFO

**S1 Table.** BChron output table.

## References

1. Brown B. The Hell Creek beds of the Upper Cretaceous of Montana: their relation to contiguous deposits, with faunal and floral lists, and a discussion of their correlation. Bulletin of the American Museum of Natural History. 1907; 23(33).

2. Archibald JD. A Study of Mammalia and Geology across the Cretaceous-Tertiary Boundary in Garfield County, Montana: University of California Publication in Geological Sciences. 1982; 122: 286 p.

3. Archibald JD, Butler RF, Lindsay EH, Clemens, WA, Dingus L. Upper Cretaceous–Paleocene biostratigraphy and magnetostratigraphy, Hell Creek and Tullock Formations, northeastern Montana: Geology. 1982; 10:153–159. doi:10.1130/0091-7613

4. Swisher CC, Dingus L, Butler RF. Ar/Ar dating and magnetostratigraphic correlation of the terrestrial Cretaceous-Paleogene boundary and Puercan mammal age, Hell Creek–Tullock Formations, eastern Montana: Canadian Journal of Earth Sciences. 1993; 30:1981–1996. doi:10.1139/e93-174.

5. Clemens WA. Evolution of the mammalian fauna across the Cretaceous-Tertiary boundary in northeastern Montana and other areas of the western interior. In: Hartman JH, Johnson KR, Nichols DJ, editors. The Hell Creek Formation and the Cretaceous-Tertiary Boundary in the Northern Great Plains: An Integrated Continental Record of the End of the Cretaceous: Geological Society of America Special Paper 361. 2002; p. 217–245.

6. Johnson KR, Nichols DJ, Hartman JH. Hell Creek Formation: A 2001 synthesis. In: Hartman JH, Johnson KR, Nichols DJ, editors. The Hell Creek Formation and the Cretaceous-Tertiary Boundary in the Northern Great Plains: An Integrated Continental Record of the End of the Cretaceous: Geological Society of America Special Paper 361. 2002; p. 503–510.

7. Murphy E, Hoganson J, Johnson K. Lithostratigraphy of the Hell Creek Formation in North Dakota. In: Hartman JH, Johnson KR, Nichols DJ, editors. The Hell Creek Formation and the Cretaceous-Tertiary Boundary in the Northern Great Plains: An Integrated Continental Record of the End of the Cretaceous: Geological Society of America Special Paper 361. 2002; p. 9–34.

8. Pearson, D.A., Schaefer, T., Johnson, K.R., Nichols, D.J., and Hunter, J.P., 2002, Vertebrate biostratigraphy of the Hell Creek Formation in southwestern North Dakota and northwestern South Dakota. In: Hartman JH, Johnson KR, Nichols DJ, editors. The Hell Creek Formation and the Cretaceous-Tertiary Boundary in the Northern Great Plains: An Integrated Continental Record of the End of the Cretaceous: Geological Society of America Special Paper 361. 2002; p. 145–167.

9. Wilson GP. Mammalian faunal dynamics during the last 1.8 million years of the Cretaceous in Garfield County, Montana: Journal of Mammalian Evolution. 2005; 12, 53–76, doi:10.1007/s10914-005-6943-4.

10. Horner, JR, Goodwin MB, Myhrvold N. Dinosaur Census Reveals Abundant Tyrannosaurus and Rare Ontogenetic Stages in the Upper Cretaceous Hell Creek Formation (Maastrichtian), Montana, USA. PLoS ONE. 2011; 6(2): e16574. doi:10.1371/journal.pone.0016574.

11. Hartman JH, Butler RD, Weiler MW, Schumaker KK. Context, naming, and formal designation of the Cretaceous Hell Creek Formation lectostratotype, Garfield County, Montana. In Wilson GP, Clemens WA, Horner JR, Hartman JH, editors. Through the End of the Cretaceous in the Type Locality of the Hell Creek Formation in Montana and Adjacent Areas. Geological Society of America. 2014. pp. 89–121. doi:10.1130/2014.2503(02).

12. Scannella, J.B., Fowler, D.W., Goodwin, M.B., Horner, J.R. Evolutionary trends in *Triceratops* from the Hell Creek Formation, Montana. PNAS. 2014; 111(28):10245–10250.

13. Wilson GP. Mammalian extinction, survival, and recovery dynamics across the Cretaceous-Paleogene boundary in northeastern Montana, USA. In: Wilson GP, Clemens WA, Horner JR, Hartman JH, editors, Through the end of the Cretaceous in the type locality of the Hell Creek Formation in Montana and adjacent areas. Geological Society of America Special Papers 503. 2014. pp. 365–392.

14. Fastovsky DE, Bercovici A. The Hell Creek Formation and its contribution to the Cretaceous-Paleogene extinction: A short primer. Cretaceous Research. 2014; 57. pp. 368–390. doi: 10.1016/j.cretres.2015.07.007

15. DePalma RA, Smit J, Burnham DA, Kuiper K, Manning PL, Oleinik A, Larson P, Maurrasse FJ, Vellekoop J, Richards MA, Gurche L, Alvarez W. A seismically induced onshore surge deposit at the K-Pg boundary, North Dakota. PNAS. 2019; 116 (17): 8190–8199. doi10.1073/pnas.1817407116

16. Lyson TR, Miller IM, Bercovici A., Weissenburger K, Fuentes AJ, Clyde WC., Hagadorn JW, Butrim MJ, Johnson KR, Fleming RF, Barclay RS, Maccracken SA, Lloyd B, Wilson GP, Krause DW, Chester, SGB. Exceptional continental record of biotic recovery after the Cretaceous-Paleogene mass extinction. Science. 2019; 366, 977–983.

17. Hilton EJ, During MAD, Grande L, Alberg PE. New paddlefishes (Acipenseriformes, Polyodontidae) from the Late Cretaceous Tanis Site of the Hell Creek Formation in North Dakota, USA Journal of Paleontology. 2023; 0022-3360/23/1937-2337. doi:10.1017/jpa.2023.19

18. Atkins-Weltman KL, Simon DJ, Woodward HN, Funston GF, Snively E. A new oviraptorosaur (Dinosauria: Theropoda) from the end-Maastrichtian Hell Creek Formation of North America. PLoS ONE. 2024; 19(1): e0294901. 10.1371/journal.pone.0294901

19. LeCain R, Clyde WC, Wilson GP, Riedel J. Magnetostratigraphy of the Hell Creek and lower Fort Union Formations in northeastern Montana. In: Wilson GP, Clemens WA, Horner, JR, Hartman JH, editors. Through the End of the Cretaceous in the Type Locality of the Hell Creek Formation in Montana and Adjacent Areas: Geological Society of America Special Paper 503. 2014; p. 137–147. 10.1130/2014.2503(04)

20. Sprain CJ, Renne PR, Wilson GP, Clemens WA. High-resolution chronostratigraphy of the terrestrial Cretaceous-Paleogene transition and recovery interval in the Hell Creek region, Montana. Geological Society of America Bulletin. 2015; 127(3-4): pp. 393–409. doi:10.1130/B31076.1

21. Sprain CJ, Renne PR, Clemens WA, Wilson GP. Calibration of chron C29R: New high-precision geochronologic and paleomagnetic constraints from the Hell Creek region, Montana. Geological Society of America Bulletin. 2018; 130(9-10): 1615–1644. doi: 10.1130/B31890.1

22. Hicks JF, Johnson KR, Obradovich JD, Tauxe L, Clark D. Magnetostratigraphy and geochronology of the Hell Creek and basal Fort Union Formations of southwestern North Dakota and a recalibration of the age of the Cretaceous-Tertiary boundary. In: Hartman JH, Johnson KR, Nichols DJ, editors. The Hell Creek Formation and the Cretaceous-Tertiary Boundary in the Northern Great Plains: An Integrated Continental Record of the End of the Cretaceous: Geological Society of America Special Paper 361. 2002; pp. 35–55.

23. Wilson GP. A quantitative assessment of evolutionary and ecological change in mammalian faunas leading up to and across the Cretaceous-Tertiary boundary in northeastern Montana [Unpublished Doctoral thesis]. 2002. University of California.

24. Arens NC, Jahren AH, Kendrick DC. Carbon isotope stratigraphy and correlation of plant megafossil localities in the Hell Creek Formation of eastern Montana, USA. Geological Society of America Special Papers 503. 2014; 149–171.

25. Eberth DA, Kamo SL. First high-precision U-Pb CA-ID-TIMS age for the Battle Formation (Upper Cretaceous), Red Deer River valley, Alberta, Canada: implications for ages, correlations, and dinosaur biostratigraphy of the Scollard, Frenchman, and Hell Creek formations. Canadian Journal of Earth Sciences. 2019; 56(10): 1041–1051. doi:10.1139/cjes-2018-0098

26. Fowler D. The Hell Creek Formation, Montana: A Stratigraphic Review and Revision Based on a Sequence Stratigraphic Approach. Geosciences. 2020; 10 (11). doi:10.3390/geosciences10110435

27. Clyde WA, Ramezani JA, Johnson KR, Bowring SA, Jones MM. Direct high-precision U–Pb geochronology of the end-Cretaceous extinction and calibration of Paleocene astronomical timescales: Earth and Planetary Science Letters. 2016; 452: 272–280.

28. Roberts EM, Sampson SD, Deino AD, Buchwaldt R, Bowring SA, The Kaiparowits Formation: a remarkable record of Upper Cretaceous Terrestrial Ecosystems, Evolution and Tectonics in Western North America. In: Titus A, Loewen MA, editors. Advances in Late Cretaceous Western Interior Paleontology and Geology. 2013; pgs. 85–106. Indiana Press, Bloomington.

29. Beveridge TL, Roberts EM, Titus AL. Volcaniclastic member of the richly fossiliferous Kaiparowits Formations reveals new insights for regional correlation and tectonics of the southern Cordilleran Foreland basin during the latest Campanian. Cretaceous Research. 2020; 114: 104527.

30. Beveridge TL, Roberts EM, Ramezani J, Titus AL, Eaton JG, Irmis RB, Sertich JJW. Refined geochronology and revised stratigraphic nomenclature of the Upper Cretaceous Wahweap Formation, Utah, U.S.A. and the age of early Campanian vertebrates from southern Laramidia. Palaeogeography, Palaeoclimatology, Palaeoecology. 2022; 591:110876.

31. Ramezani J, Beveridge TL, Rogers RR, Eberth DA, Roberts EM. Calibrating the zenith of dinosaur diversity in the Campanian of the Western Interior Basin by CA-ID-TIMS U-Pb Geochronology. Scientific Reports. 2022; 12:16026.

32. Eberth DA, Evans DC, Ramezani J, Kamo, SL, Brown CM, Currie PJ., Braman, DR. Calibrating geologic strata, dinosaurs, and other fossils at Dinosaur Provincial Park (Alberta, Canada) using a new CA-ID-TIMS U–Pb geochronology. Canadian Journal of Earth Sciences. 2023; 60 (12). 10.1139/cjes-2023-0037

33. Rogers RR, Eberth DA, Ramezani J. The “Judith River–Belly River problem” revisited (Mon-tana-Alberta-Saskatchewan): New perspectives on the correlation of Campanian dinosaur bearing strata based on a revised stratigraphic model updated with CA-ID-TIMS U-Pb geochronology. Geological Society of America Bulletin. 2023; 136: 1221–1237. 10.1130/B36999.1

34. Rogers RR, Horner JR, Ramezani R, Roberts ER, Varricchio, DJ. Updating the Upper Cretaceous (Campanian) Two Medicine Formation of Montana: Lithostratigraphic revisions, new CA-ID-TIMS U-Pb ages, and a calibrated framework for dinosaur occurrences. 2025; 137 (1/2): 315–340. 10.1130/B37498.1

35. Singer BS, Jicha BR, Sawyer DA, Walaszczyk I, Landman N, Sageman BB, McKinney KC. A ^40^Ar/^39^Ar and U-Pb timescale for the Cretaceous Western Interior Basin, North America. Geological Society, London, Special Publications. 2023; 544: 367–391. 10.1144/SP544-2023-7

36. Hartman JH. Hell Creek Formation and the early picking of the Cretaceous-Tertiary boundary in the Williston Basin. In: Hartman JH, Johnson KR, Nichols DJ, editors. The Hell Creek Formation and the Cretaceous-Tertiary Boundary in the Northern Great Plains: An Integrated Continental Record of the End of the Cretaceous: Geological Society of America Special Paper 361. 2002; pp.1–7.

37. Kaskes P. Unearthing the background of Naturalis Tyrannosaurus rex: taphonomy, stratigraphy and paleoenvironment [Unpublished Masters Thesis]. 2016. University of Amsterdam.

38. Sager M. Will the Public Ever Get to See the “Duelling Dinosaurs”? Smithsonian Magazine. 2017; July. https://www.smithsonianmag.com/science-nature/public-ever-see-dueling-dinosaurs-180963676/

39. Cobban WA, Reeside JB. Correlation of the Cretaceous formations of the western interior of the United States. Geological Society of America Bulletin. 1952; 63(10):1011–1044.

40. Frye CI. Stratigraphy of the Hell Creek Formation in North Dakota. North Dakota Geological Survey Bulletin. 1969; 54, 1–65.

41. Hares CJ. Geology and lignite resources of the Marmarth Field, southwestern North Dakota: U.S. Geological Survey Bulletin. 1928; 775, 110 p.

42. Fastovsky DE, Bercovici A. The Hell Creek Formation and its contribution to the Cretaceous– Paleogene extinction: A short primer. Cretaceous Research. 2016; 57: 368–390.

43. Fuentes AJ, Clyde WC, Weissenburger K, Bercovici A, Lyson TR, Miller IM, Ramezani J, Isakson V, Schmitz MD, Johnson KR. Constructing a time scale of biotic recovery across the Cretaceous-Paleogene boundary, Corral Bluffs, Denver Basin, Colorado, U.S.A. Rocky Mountain Geology. 2019; 54:133–153.

44. Lawton TF, Talling PJ, Hobbs RS, Trexler JH, Weiss MP, Burbank DW. Structure and stratigraphy of Upper Cretaceous and Paleogene strata (North Horn Formation), Eastern San Pitch Mountains, Utah—Sedimentation at the front of the Sevier orogenic front. U.S. Geological Survey Bulletin 1993; 1787-II. 31p.

45. Lozinsky RP, Hunt AP, Wolberg DL, Lucas SG. Late Cretaceous (Lancian) dinosaurs from the McRae Formation, Sierra County, New Mexico. New Mexico Geology. 1984; 6:72–77.

46. Hunt AP, Lucas SG. Stratigraphy, paleontology and age of the Fruitland and Kirtland Formations (Upper Cretaceous), San Juan Basin, New Mexico. In: New Mexico Geological Society Guidebook, 43rd Field Conference, San Juan Basin IV. 1992. pp. 218–239.

47. Leslie CE, Peppe, DJ, Williamson TE, Heizler M, Jackson M, Atchley SC, Nordt L, Standhardt B. Revised age constraints for Late Cretaceous to early Paleocene terrestrial strata from the Dawson Creek section, Big Bend National Park, west Texas. Geological Society of America Bulletin. 2018; 130 (7-8): 1143–1163. 10.1130/B31785.1

48. Alvarez LW, Alvarez W, Asaro F, Michel HV. Extraterrestrial Cause for the Cretaceous-Tertiary Extinction. Science. 1980; 208(4448):1095–1108. doi:10.1126/science.208.4448.1095

49. Smit J, Kaars SVD. Terminal Cretaceous Extinctions in the Hell Creek Area, Montana: Compatible with Catastrophic Extinction. Science. 1984; 223, 1177–1179.

50. Johnson KR. Leaf-fossil evidence for extensive floral extinction at the Cretaceous-Tertiary boundary, North Dakota, USA: Cretaceous Research. 1992; 13 (1): 91–117.

51. Renne PR, Deino AL, Hilgen FJ, Kuiper KF, Mark DF, Smit J. Time Scales of Critical events Around the Cretaceous-Paleogene Boundary. Science. 2013; 339 (6120): 684–687.

52. Deibel PKW, Wilson Mantilla GP, Strömberg CAE. Plant taxonomic turnover and diversity across the Cretaceous/Paleogene boundary in northeastern Montana. Paleobiology. 2024; 50: 608– 636. doi:10.1017/pab.2024.22

53. Lofgren DL. The Bug Creek Problem and the Cretaceous-Tertiary Transition at McGuire Creek, Montana. University of California Publ. Geol. Sci. 1995; 140: 204p.

54. Scholz H, Hartman JS. Paleoenvironmental reconstruction of the Upper Cretaceous Hell Creek Formation of the Williston Basin, Montana, USA: Implications from the quantitative analysis of Unionoid bivalve taxonomic diversity and morphologic disparity. PALAIOS. 2007; 22(1): 24–34. doi: 10.2110/palo.2005.p05-059r

55. Clemens WA, Hartman JH. From Tyrannosaurus rex to asteroid impact: Early studies (1901-1980) of the Hell Creek Formation in its type area. In: Wilson GP, Clemens WA, Horner JR, Hartman JH, editors. Through the End of the Cretaceous in the Type Locality of the Hell Creek Formation in Montana and Adjacent Areas. Geological Society of America. 2014. pp. 89–121.

56. Pearson DA, Schaefer T, Johnson KR, Nichols DJ. Palynologically calibrate vertebrate record from North Dakota consistent with abrupt dinosaur extinction at the Cretaceous-Tertiary boundary. Geology. 2001; 29(1): 39–42.

57. Lerbekmo JF. Glacioeustatic sea level fall marking the base of supercycle TA-1 at 66.5 Ma recorded by the kaolinization of the Whitemud Formation and the Colgate Member of the Fox Hills Formation. Marine and Petroleum Geology. 2009; 26(7): 1299–1303. doi:10.1016/j.marpetgeo.2008.08.001

58. Sprain CJ, Feinberg JM, Renne PR, Jackson M. Importance of titanohematite in detrital remanent magnetizations of strata spanning the Cretaceous-Paleogene boundary, Hell Creek region, Montana: Geochemistry, Geophysics, Geosystems. 2016; 17 (3): 660–678. doi: 10.1002/2015GC006191.

59. Lurcock PC, Wilson GS. PuffinPlot: A versatile, user-friendly program for paleomagnetic analysis. Geochemistry, Geophysics, Geosystems. 2012; 13 (6). 10.1029/2012GC004098

60. Kirschvink JL. The least-squares line and plane and the analysis of palaeomagnetic data: Geophysical Journal International. 1980; 62: 699–718, 10.1111/j.1365-246X.1980.tb02601.x.

61. Fisher RA. Dispersion on a sphere: Proceedings of the Royal Society of London, Series A, Mathematical and Physical Sciences. 1953; 217: 295–305.

62. Hoke GD, Schmitz M, Bowring SA. An ultrasonic method for isolating nonclay components from clay-rich material. Geochemistry, Geophysics, Geosystems. 2014; 15(2): 492–498.

63. Mattinson JM. Zircon U-Pb chemical abrasion (“CA-TIMS”) method: Combined annealing and multi-step partial dissolution analysis for improved precision and accuracy of zircon ages. Chemical Geology. 2005; 220(1-2): 47–66. doi: 10.1016/j.chemgeo.2005.03.011

64. Condon DJ, Schoene B, McLean NM, Bowring SA, Parrish, RR. Metrology and traceability of U-Pb isotope dilution geochronology (EARTHTIME Tracer Calibration Part I). Geochimica et Cosmochimica Acta. 2015; 164: 464–480. 10.1016/j.gca.2015.05.026

65. McLean NM, Condon DJ, Schoene B & Bowring SA. Evaluating uncertainties in the calibration of isotopic reference materials and multi-element isotopic tracers (EARTHTIME Tracer Calibration Part II). Geochimica et Cosmochimica Acta. 2015; 164: 481–501. 10.1016/j.gca.2015.02.040

66. Krogh TE. Low-contamination method for hydrothermal decomposition of zircon and extraction of U and Pb for isotopic age determinations. Geochimica et Cosmochimica Acta 37: 485–494. 10.1016/0016-7037(73)90213-5

67. Gerstenberger H, Hasse G. A highly effective emitter substance for mass spectrometric Pb isotope ratio determinations. Chemical Geology. 1997; 136 (3–4): 309–312. 10.1016/S0009-2541(96)00033-2

68. Bowring JF, McLean NM, Bowring SA. Engineering cyber infrastructure for U-Pb geochronology: Tripoli and U-Pb Redux. Geochemistry, Geophysics, Geosystems. 2011; 12 (6): Q0AA19. doi.org/10.1029/2010gc003479

69. McLean, N. M., Bowring, J. F. & Bowring, S. A. An algorithm for U-Pb isotope dilution data reduction and uncertainty propagation. Geochemistry, Geophysics, Geosystems. 2011; 12 (6): Q0AA18. doi.org/10.1029/2010gc003478

70. Machlus ML, Ramezani J, Bowring SA, Hemming SR A strategy for cross-calibrating U-Pb chronology and astrochronology of sedimentary sequences: an example from the Green River Formation, Wyoming, USA. Earth and Planetary Science Letters. 2015; 413:70–78. 10.1016/j.epsl.2014.12.009

71. Hiess J, Condon DJ, McLean N, Noble SR. ^238^U/^235^U systematics in terrestrial uranium-bearing minerals. Science. 2012; 335:1610–1614. 10.1126/science.1215507

72. Mattinson JM. Analysis of the relative decay constants of ^235^U and ^238^U by multi-step CA-TIMS measurements of closed-system natural zircon samples. Chemical Geology. 2010; 275: 186–198. 10.1016/j.chemgeo.2010.05.007

73. Schoene B, Crowley JL, Condon DJ, Schmitz MD, Bowring SA. Reassessing the uranium decay constants for geochronology using ID-TIMS U-Pb data. Geochimica et Cosmochimica Acta. 2006; 70: 426–445. 10.1016/j.gca.2005.09.007

74. Jaffey AH, Flynn KF, Glendenin LE, Bentley WC, Essling AM. Precision measurement of half-lives and specific activities of ^235^U and ^238^U. Physical Review C. 1971; 4:1889–1906. 10.1103/PhysRevC.4.1889

75. Nasdala L, Corfu F, Schoene B, Tapster SR, Wall, CJ, Schmitz MD, et al. GZ7 and GZ8—two zircon reference materials for SIMS U-Pb geochronology. Geostandards and Geoanalytical Research. 2018; 42:431–457. 10.1111/ggr.12239

76. Haslett J, Parnell AA. simple monotone process with application to radiocarbon-dated depth chronologies. Journal of the Royal Statistics Society Applied Statistics C. 2008; 57:399–418. 10.1111/j.1467-9876.2008.00623.x

77. Parnell AC, Haslett J, Allen JRM, Buck CE, Huntley B. A flexible approach to assessing synchroneity of past events using Bayesian reconstructions of sedimentation history. Quaternary Science Reviews. 2008; 27:1872–1885. 10.1016/j.quascirev.2008.07.009

78. Ogg JG. Chapter 5 – Geomagnetic Polarity Time Scale. Felix M. Gradstein, James G. Ogg, Mark D. Schmitz, Gabi M. Ogg, editors. Geologic Time Scale 2020, Elsevier. 2020. Pgs.159–192. 10.1016/B978-0-12-824360-2.00005-X.

79. Thom WT, Dobbin CE. Stratigraphy of Cretaceous-Eocene transition beds in eastern Montana and the Dakotas. Geological Society of America Bulletin. 1924; 35: 481–506.

80. Nichols DJ, Brown JL. Palynostratigraphy of the Tullock Member (lower Paleocene) of the Fort Union Formation in the Powder River basin, Montana and Wyoming. U.S. Geological Survey Bulletin, 1917-F. 1992: 35 pp.

81. Pocknall DT, Nichols DJ. Palynology of Coal Zones of the Tongue River Member (upper Paleocene) of the Fort Union Formation, Powder River Basin, Montana and Wyoming. Issue 32 of Contributions series, American Association of Stratigraphic Palynologists. 1996. American Association of Stratigraphic Palynologists Foundation. 58 p.

82. Nichols DJ. Palynology and palynostratigraphy of the Hell Creek Formation in North Dakota: A microfossil record of plants at the end of Cretaceous time. In: Hartman JH, Johnson KR, Nichols DJ, editors. The Hell Creek Formation and the Cretaceous-Tertiary Boundary in the Northern Great Plains: An Integrated Continental Record of the End of the Cretaceous: Geological Society of America Special Paper 361. 2002; p. 393–456.

83. Nichols DJ. Palynostratigraphic framework for age determination and correlation of the nonmarine lower Cenozoic of the Rocky Mountains and Great Plains region. In: Raynolds RG, Flores RM, editors. Cenozoic systems of the Rocky Mountain Region. Denver, Colorado. Rocky Mountain Section of the Society for Sedimentary Geology (SEPM) 2003. pp. 107e134.

84. Nichols DJ, Ott HL. Neotypes for Paleocene species in the *Momipites-Caryapollenites* pollen lineage. Palynology. 2006; 30 (1): 33–41.

85. Bercovici A, Pearson D, Nichols D, Wood J. Biostratigraphy of selected K/T boundary sections in southwestern North Dakota, USA: Toward a refinement of palynological identification criteria. Cretaceous Research. 2009; 30: 632–658.

86. Braman, DR, Sweet AR. Biostratigraphically useful Late Cretaceous–Paleocene terrestrial palynomorphs from the Canadian Western Interior sedimentary basin. Palynology. 2012; 36:8–35.

87. Braman DR. Terrestrial palynostratigraphy of the Upper Cretaceous (Santonian) to lowermost Paleocene of southern Alberta, Canada. Palynology. 2018; 42: 102–147.

88. Nichols DJ, Johnson KR. Palynology and microstratigraphy of Cretaceous-Tertiary boundary sections in southwester North Dakota. In: Hartman JH, Johnson KR, Nichols DJ, editors. The Hell Creek Formation and the Cretaceous-Tertiary Boundary in the Northern Great Plains: An Integrated Continental Record of the End of the Cretaceous: Geological Society of America Special Paper 361. 2002; pp. 95–143.

89. Nadon GC. The genesis and recognition of anastomosed fluvial deposits: data from the St. Mary River Formation, Southwestern Alberta, Canada. Journal of Sedimentary Research. 1994; B64 (4): 451–463.

90. Connor, CW. The Lance Formation–Petrography and Stratigraphy, Powder River Basin and Nearby Basins, Wyoming and Montana. US Geological Survey Bulletin 1917-I: Evolution of the sedimentary basins–Powder River Basin. 1980; 117p.

91. Smith RL. Ash flows. Geological Society of America Bulletin. 1960; 71: 795–841.

92. Smedes HW. Geology and igneous petrology of the northern Elkhorn mountains. United States Geological Survey Professional Paper 510. 1966: 116p.

93. Thomas RG, Eberth DA, Deino AL, Robinson D. Composition, radioisotopic ages, and potential significance of an altered volcanic ash (bentonite) from the Upper Cretaceous Judith River Formation, Dinosaur Provincial Park, southern Alberta, Canada. Cretaceous Research. 1990; 11: 125–162. 10.1016/s0195-6671(05)80030-8

94. Rogers RR, Swisher CC, Horner JR. ^40^Ar/^39^Ar age and correlation of the nonmarine Two Medicine Formation (Upper Cretaceous), northwestern Montana, U.S.A. Canadian Journal of Earth Sciences. 1993; 30:1066–1075. 10.1139/e93-090

95. Roberts EM, Hendrix MS. Taphonomy of a petrified forest in the Two Medicine formation (Campanian) northwest Montana: Implications for palinspastic restoration of the Boulder batholith and Elkhorn Mountains Volcanics. PALAIOS. 2000; 15(5): 476–482.

